# Cocaine Triggers Glial-Mediated Synaptogenesis

**DOI:** 10.1101/2020.01.20.896233

**Authors:** Junshi Wang, King-Lun Li, Avani Shukla, Ania Beroun, Masago Ishikawa, Xiaojie Huang, Yao Wang, Yao Q. Wang, Noah D. Bastola, Hugh H. Huang, Lily E. Kramer, Terry Chao, Yanhua H. Huang, Susan R. Sesack, Eric J. Nestler, Oliver M. Schlüter, Yan Dong

## Abstract

Synaptogenesis is essential in forming new neurocircuits during development, and this is mediated in part by astrocyte-released thrombospondins (TSPs) and activation of their neuronal receptor, α2δ-1. Here, we show that this developmental synaptogenic mechanism is utilized during cocaine experience to induce spinogenesis and the generation of AMPA receptor-silent glutamatergic synapses in the adult nucleus accumbens (NAc). Specifically, cocaine administration activates NAc astrocytes, and preventing this activation blocks cocaine-induced generation of silent synapses. Furthermore, knockout of TSP2, or pharmacological inhibition or viral-mediated knockdown of α2δ-1, prevents cocaine-induced generation of silent synapses. Moreover, disrupting TSP2-α2δ-1-mediated spinogenesis and silent synapse generation in the NAc occludes cue-induced cocaine seeking after withdrawal from cocaine self-administration and cue-induced reinstatement of cocaine seeking after drug extinction. These results establish that silent synapses are generated by an astrocyte-mediated synaptogenic mechanism in response to cocaine experience and embed critical cue-associated memory traces that promote cocaine relapse.

## Introduction

Experiencing cocaine and other drugs of abuse remodels glutamatergic transmission in the NAc (NAc), which underlies key circuit mechanisms that promote drug seeking and relapse after withdrawal (Kalivas, 2009; Wolf, 2010). This process is partially mediated by AMPA receptor (AMPAR)-silent glutamatergic synapses, which are generated in the NAc after repeated exposure to cocaine, and become gradually ‘un-silenced’ during prolonged drug withdrawal by recruiting AMPARs (Dong and Nestler, 2014). The resulting circuit remodeling promotes cue-induced cocaine seeking, a behavioral readout of cue-associative cocaine memory (Brown et al., 2011; Graziane et al., 2016; Huang et al., 2009; Lee et al., 2013; Ma et al., 2014). Upon cue re-exposure after cocaine withdrawal, these synapses are temporarily re-silenced, contributing to temporary destabilization of cocaine memories, and experimentally locking these synapses in the AMPAR-silent state reduces subsequent cue-induced cocaine seeking (Wright et al., 2019). As such, NAc silent synapses may stand as a unique set of neuronal substrates embedding memory traces that promote cocaine relapse. Although it has been hypothesized that these NAc silent synapses are generated de novo by cocaine experience (Dong and Nestler, 2014), the cellular mechanisms that mediate such potentially large-scale synaptogenesis in the adult brain remain incompletely understood.

Virtually all current efforts at understanding cocaine-induced generation of silent synapses have focused on neuronal mechanisms. First, silent synapse generation in response to cocaine requires activation of CREB, a transcription factor critically implicated in synaptogenesis and circuit formation during development (Brown et al., 2011; Lonze and Ginty, 2002). Second, cocaine-generated silent synapses are enriched in GluN2B-containing NMDA receptors (NMDARs), a signature feature of nascent excitatory synapses during development (Brown et al., 2011; Cull-Candy and Leszkiewicz, 2004; Graziane et al., 2016; Huang et al., 2009). Third, cocaine-induced generation of silent synapses is accompanied by an increase in the density of dendritic spines, postsynaptic structures of glutamatergic synapses, and these effects are mediated in part by small G proteins (Cahill et al., 2016; Graziane et al., 2016; Wright et al., 2019; Yuste and Denk, 1995). These features support our hypothesis that cocaine experience activates and utilizes developmental synaptogenic mechanisms in the adult brain to generate nascent silent synapses.

Despite the importance of neuronal mechanisms, work to date has not considered glial mechanisms in cocaine action which, in the developing brain, are also crucial for generating AMPAR-silent glutamatergic synapses. Prominent among such glial-generated signals are astrocyte-secreted thrombospondins (TSP) 1 and 2 (encoded by the Thbs1 and Thbs2 genes), which are both sufficient and necessary for new synapse formation (Clarke and Barres, 2013). By activating their neuronal receptor, α2δ-1, TSP1/2 induce pre-versus postsynaptic differentiation and clustering of synaptic proteins, resulting in NMDAR-enriched, AMPAR-silent glutamatergic synapses (Christopherson et al., 2005; Eroglu et al., 2009). After development, the astrocytic TSP1/2-α2δ-1 signaling substantially declines, which may contribute to the relatively low levels of silent synapses and reduced synaptogenesis in the adult brain (Christopherson et al., 2005; Hoffman et al., 1994; Iruela-Arispe et al., 1993).

In the present study, we show that cocaine administration acutely activates astrocytes in NAc shell (NAcSh) slices, and that preventing astrocyte activation in vivo prevents cocaine-induced generation of silent synapses. Furthermore, disrupting TSP2-α2δ-1 signaling abolishes cocaine-induced increases in the density of dendritic spines as well as the generation of silent synapses in the NAcSh. Moreover, preventing TSP2-α2δ-1-mediated synaptogenesis in the NAcSh during cocaine self-administration attenuates both cue-induced cocaine seeking after drug withdrawal and undermines cue-induced reinstatement of cocaine seeking after drug extinction. Thus, cocaine experience activates a developmental, astrocyte-mediated synaptogenic mechanism to generate nascent, AMPAR-silent synapses, which may embed critical memory traces that promote cue-induced drug relapse.

## Results

### Cocaine activation of astrocytes

Repeated exposure to cocaine generates AMPAR-silent glutamatergic synapses in the NAcSh (Brown et al., 2011; Huang et al., 2009). These synapses share several core features with nascent glutamatergic synapses during brain development (Dong and Nestler, 2014; Huang et al., 2015b), during which astrocyte-controlled synaptogenesis is highly active (Allen and Eroglu, 2017; Chung et al., 2015). Here, we tested whether cocaine-generated silent synapses in adulthood are also regulated by astrocyte-mediated mechanisms.

In the NAcSh from adult rats (20-week old), our electron microscopy studies detected excitatory synapses ensheathed by astrocytes, indicating an anatomical basis for astrocytic regulation of NAcSh synapses (**Fig. 1A**). Astrocytes respond to neuronal activation and to certain external stimuli with increases in intracellular Ca^2+^ levels (Volterra and Meldolesi, 2005). To monitor such Ca^2+^-mediated astrocytic activations, we crossed the Aldh1/1-Cre/ERT2 mouse line with the ROSA26-Lck-GCaMP6f-flox line to generate the mice in which astrocyte-specific expression of GCaMP6f can be induced by tamoxifen (Srinivasan et al., 2016). In these mice, the Lck moiety tethers the expressed GCaMP6f to fine processes of astrocytes, resulting in unique bushy expression patterns (Madisen et al., 2015; Srinivasan et al., 2016). Through immunohistochemistry and confocal microscopy, we imaged Lck-GCaMP6f-GFP in the NAcSh and observed such bushy expression patterns, which are typical of astrocytes but overlapped minimally with NeuN-labeled neurons (**Figs. 1B-D; S1A,B**). We prepared live NAcSh slices from these mice (8-16-week old) after tamoxifen induction and imaged GCaMP6f-mediated fluorescent signals (representing astrocytic Ca^2+^ increases) before and during perfusion of cocaine (20 μM) (**Fig. 1E**). Spontaneous Ca^2+^ increases were detected within many small, relatively well-defined locations (diameter ∼5-15 μm) throughout the NAcSh slice. These Ca^2+^-mediated events may represent individually activated astrocytes or independently activated domains within the same astrocytes. Analyzing the GCaMP6f-mediated transients of these individual Ca^2+^-mediated events (**Fig. 1K**; also see **Methods**) revealed that perfusion of cocaine increased the number of locations that exhibited Ca^2+^-mediated events within the same slices (**Fig. 1L**), indicating that some dormant astrocytes or dormant domains within the same astrocytes were activated by cocaine. Furthermore, perfusion of cocaine also increased the total number of Ca^2+^-mediated events (**Fig. 1M**), and this effect was likely attributable to both the increased number of activation locations and increased activation frequencies of the same locations (**Fig. 1N**). Consequently, the overall astrocytic Ca^2+^ increases, estimated by the cumulative areas swept by GCaMP6f-mediated transients of all Ca^2+^-mediated events, were also increased during perfusion of cocaine (**Fig. 1O**). Meanwhile, the mean and maximum amplitudes of GCaMP6f-mediated transients of individual Ca^2+^-mediated events were similar during perfusion of control artificial cerebrospinal fluid (aCSF) and cocaine (**Fig. S1C,D**). This observation not only indicated minimal fluorescence bleaching during aCSF versus cocaine perfusion, but also suggested that the overall increased astrocytic Ca^2+^ activation was not attributable to increased activation intensity of individual Ca^2+^ events.

**Figure 1.**
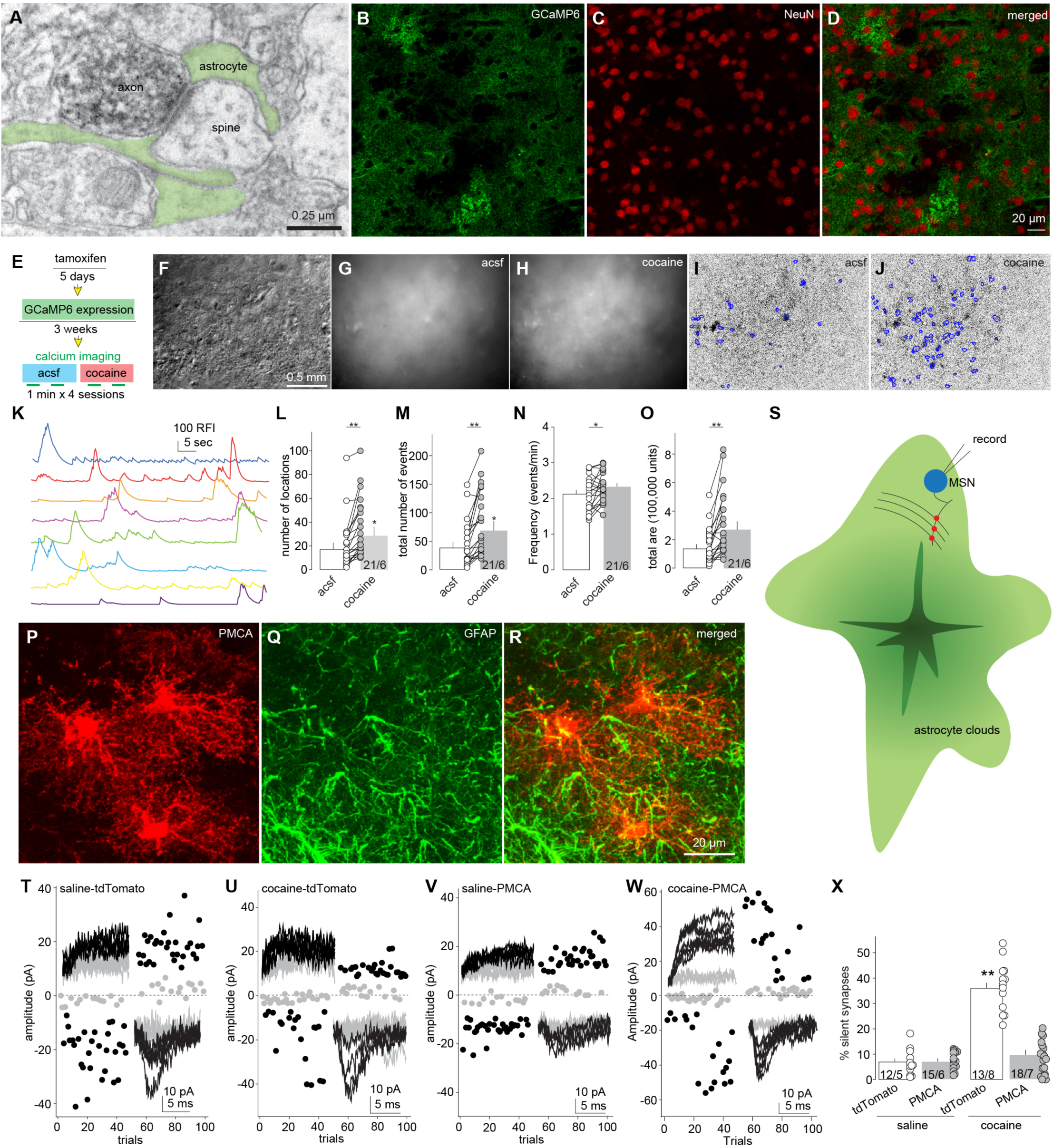
Cocaine-induced generation of silent synapses requires activation of astrocytes. **A** Electronic microscopic image showing a representative NAc excitatory synapse exhibiting immunoperoxidase labeling for vGlut1 in the axon and ensheathment by astrocytes (green shading). **B**-**D** Example confocal images of an NAcSh slice from a Aldh1/1-Cre/Ert2 x Rosa26-Lck-GCaMP6f-flox mouse showing astrocyte-like, bushy distributions of GCaMP6f-GFP signals (**B**), NeuN-stained MSNs (**C**), and minimal co-expression of GCaMP6f and NeuN (**D**). **E** Experimental design illustrating that after astrocyte-selective expression of GCaMP6f NAcSh slices were prepared and sequentially perfused by aCSF (control) and cocaine for calcium imaging of astrocytes. **F**-**H** Examples of unprocessed NAcSh images through DIC channel (F) and 1-min stacked images through GFP channel during perfusions of aCSF (**G**) and cocaine (**H**). **I**,**J** Extracted GCaMP6f signals of the same slice during perfusion of aCSF (**I**) or cocaine (**J**). **K** Calcium transients from 8 example locations where GCaMP6f activities were detected. **L**-**O** Summaries showing that acute perfusion of cocaine increased the number of locations where GCaMP6f signals were detected (t_1,20_= 4.73, p < 0.01; **L**), the number of total GCaMP6f-mediated events over the 1-min imaging period (t_1,20_= 4.98, p < 0.01; **M**), the number of events per min (t_1,20_= 2.36, p = 0.03; **N**), and the total area swept by calcium transients (t_1,20_= 4.44, p < 0.01;**O**). **P-R** Example confocal images of an NAcSh slice with GfaABC_1_D AAV-mediated expression of hPMCA2w/b-mCherry, in which mCherry signals depicted bushy, astrocytic morphologies (**P**), and the GFAP staining (Green) of the same slice revealed astrocytic skeletons (**Q**) that overlapped with the hPMCA2w/b-mCherry signals (**R**). **S** Diagram showing selective recordings of MSNs that were within the bushy matrix of astrocytes that expressed hPMCA2w/b-mCherry. **T**-**W** Example trials of the minimal stimulation assays of NAcSh MSNs from saline (**T**)- or cocaine-trained (**U**) rats with NAcSh astrocytic expression of tdTomato, or saline (**V**) or cocaine-trained (**W**) rats with NAcSh astrocytic expression of hPMCA2w/b-mCherry. Insets showing example EPSCs at +40 or −70 mV from the minimal stimulation assay. **X** Summary showing that astrocytic expression of hPMCA2w/b did not affect the basal % silent synapses in the NAcSh in saline-trained rats, but prevented generation of silent synapses in cocaine-trained rats (F_1,22_ = 57.83, p < 0.01, animal-based two-way ANOVA; p < 0.01 saline-tdTomato vs. cocaine-tdTomato, p = 1 saline-tdTomato vs. saline-PMCA, p<0.01 cocaine-tdTomato vs. cocaine-PMCA, p = 1 saline-PMCA vs. cocaine-PMCA, Bonferroni posttest). *, p < 0.05’ **, p < 0.01.

If occurring in vivo, cocaine-associated activation of NAcSh astrocytes may trigger release of synaptogenic factors and, if silent synapses in cocaine-trained animals are nascent synapses generated through the astrocyte-mediated synaptogenic processes, preventing the activation of NAcSh astrocytes should prevent cocaine-induced generation of silent synapses. To test this scenario, we used a modified version of the plasma membrane Ca^2+^ ATPase, namely hPMCA2w/b. Virally expressed hPMCA2w/b constitutively extrudes cytosolic Ca^2+^ and reduces Ca^2+^ elevations, and has been shown to compromise Ca^2+^-dependent signaling when expressed in striatal astrocytes (Yu et al., 2018). To deliver hPMCA2w/b into NAc astrocytes in vivo, we employed an AAV2/5 vector with the GfaABC_1_D promotor, which enables superior astrocyte-specific expression compared to other promotor options (Shigetomi et al., 2013; Yu et al., 2018). In the NAcSh, GfaABC_1_D-mediated AAV expression of hPMCA2w/b exhibited high astrocyte-specificity in both rats (**Figs. 1P-R; S1B**) and mice (**Fig. S2**). We next stereotaxically injected this hPMCA2w/b-mCherry-expressing AAV into the NAcSh of rats, using rats with injection of AAV that expressed tdTomato as controls. After 4 weeks of viral expression, we trained the rats with a 5-day cocaine self-administration procedure, and examined NAcSh silent synapses on withdrawal day 1. We performed the minimal stimulation assay within the astrocytic matrix depicted by astrocytic fluorescence, where the examined synapses were presumably under the influence of transduced astrocytes (**Figs. 1S; S1E-G**). Cocaine-trained rats with viral expression of tdTomato exhibited increased percentages of silent synapses in NAcSh medium spiny neurons (MSNs) compared to saline-trained rats, indicating that this astrocyte-specific viral expression per se did not affect cocaine-induced generation of silent synapses (**Fig. 1T-X**). In contrast, cocaine-trained rats with intra-NAcSh expression of hPMCA2w/b-mCherry exhibited comparably low % silent synapses as in saline-trained rats (**Fig. 1T-X**). Thus, preventing activation of astrocytes in the NAcSh in vivo prevents cocaine-induced generation of silent synapses.

### Role of TSP1 versus TSP2

By activating neuronal receptor α2δ-1, astrocyte-released TSPs, particularly TSP1 and TSP2, promote generation of AMPAR-silent synapses in newborn brains (Christopherson et al., 2005; Eroglu et al., 2009). While this astrocyte-mediated synaptogenic mechanism declines to low levels after development (Allen and Eroglu, 2017; Christopherson et al., 2005), it is possible that cocaine experience reactivates it to induce generation of nascent silent synapses in the NAcSh. To explore this possibility, we employed the TSP1 (ThBs1^tm1Hyn^) (Lawler et al., 1998) and TSP2 (Thbs2^tm1Bst^) (Kyriakides et al., 1998) knockout mouse lines. Similar to cocaine self-administration, repeated i.p. injections of cocaine also induce generation of silent synapses in NAcSh in both mice and rats (Brown et al., 2011; Graziane et al., 2016; Huang et al., 2009). One day after 5 days of i.p. cocaine administration, the % silent synapses in NAcSh was increased in both wild-type and TSP1 knockout mice, but not in TSP2 knockout mice, compared to saline-treated mice (**Fig. 2**). It was unfortunately not possible to determine whether cocaine alters TSP2 protein levels in the adult NAcSh due to unavailability of quality antibodies (see Discussion). Nevertheless, these results point to TSP2 as a potential astrocytic factor that mediates cocaine-induced synaptogenesis and generation of silent synapses.

**Figure 2.**
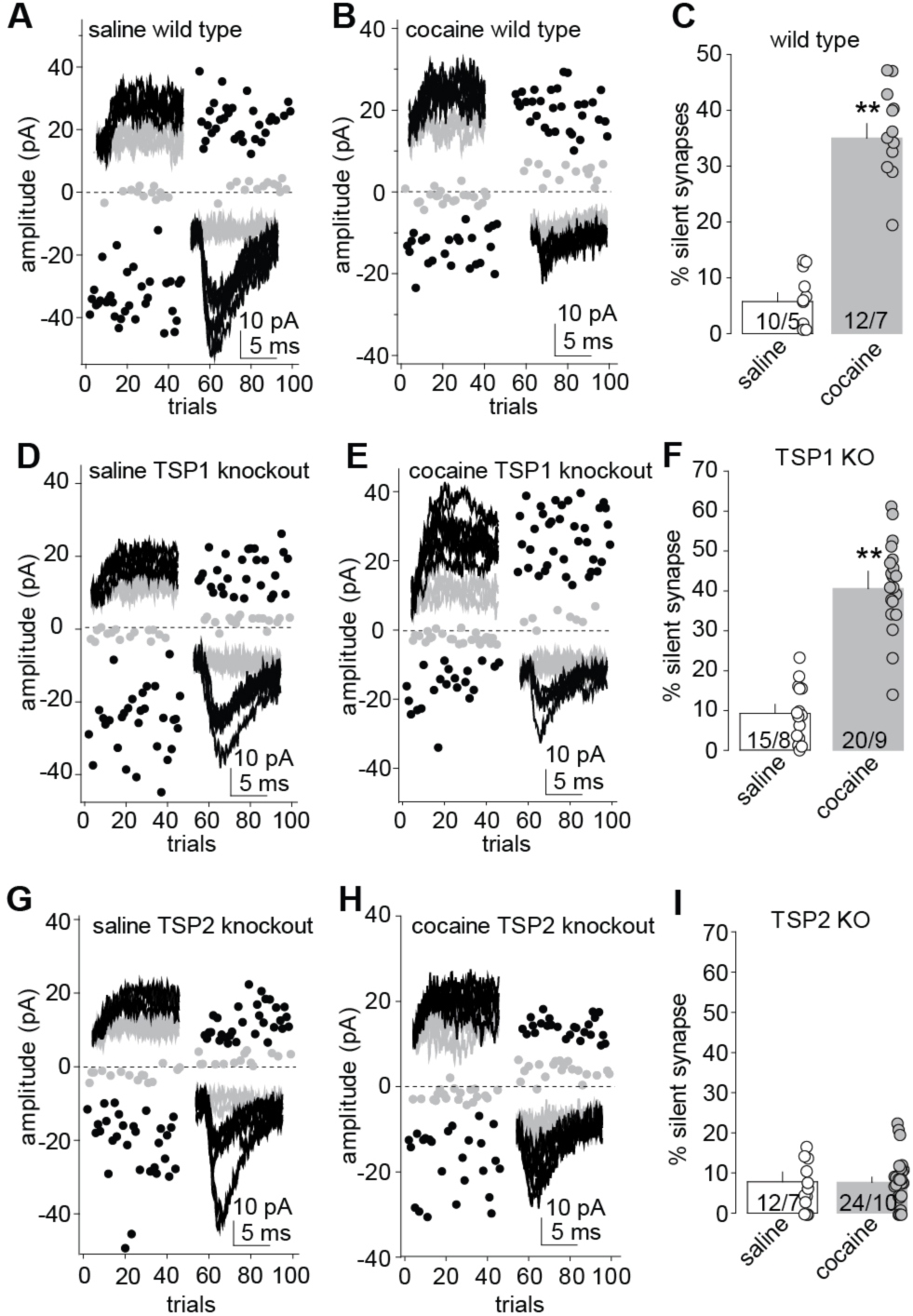
Cocaine-induced generation of silent synapses requires TSP 2. **A-C** Example trials of the minimal stimulation assay of saline- (**A**) and cocaine- (**B**) trained mice and summarized results (**C**) showing that repeated i.p. injections of cocaine generated silent synapses in NAcSh MSNs of wildtype mice (t_1,20_ = 10.18, p < 0.01, cell-based t-test; t_1,10_ = 8.62, p < 0.01, animal-based t-test). **D-F** Example trials of the minimal stimulation assay of saline (**D**) and cocaine-trained (**E**) mice and summarized results (**F**) showing that repeated i.p. injections of cocaine generated silent synapses in NAcSh MSNs of TSP 1 knockout mice (t_1,33_ = 9.68, p < 0.01, cell-based t-test; t_1,15_ = 6.10, p < 0.01, animal-based t-test). **G-I** Example trials of the minimal stimulation assay of saline (**G**) and cocaine-trained (**H**) mice and summarized results (**I**) showing that cocaine-induced generation of silent synapses was prevented in TSP 2 knockout mice (t_1,34_ = 0.87, p = 0.39, cell-based t-test; t_1,15_ = 0.07, p = 0.95, animal-based t-test). **, *p* < 0.01.

### Role of astrocytic TSP2

In addition to astrocytes, TSPs are also released from other cellular sources (i.e., neurons and endothelial cells) under physiological and pathophysiological conditions (Adams, 2001; Lin et al., 2003; Moller et al., 1996). To selectively manipulate astrocytic TSP2, we designed an shRNA (shTSP2) that targets endogenous TSP2 expression, and, through AAV2/5, drove its expression by the GfaABC_1_D promoter in a bicistronic cassette composed of the GFP coding sequence and the shTSP2 in the 3’UTR. We bilaterally injected the rats with this AAV into the NAcSh before the self-administration training (**Fig. 3A,B**). ∼4 weeks later, we co-stained GFP in transduced NAcSh slices with GFAP, an astrocyte-specific cytoskeletal protein that serves as a reliable biomarker of astrocytes in adult brain. Bushy GFP signals depicted the contour of individual astrocytes (**Fig. 3C-F**). Co-staining GFP with NeuN revealed minimal neuronal expression of shTSP2-GFP (**Fig. 3G-I**). Thus, this AAV tool allows astrocyte-specific knockdown of TSP2 in the NAcSh.

**Figure 3.**
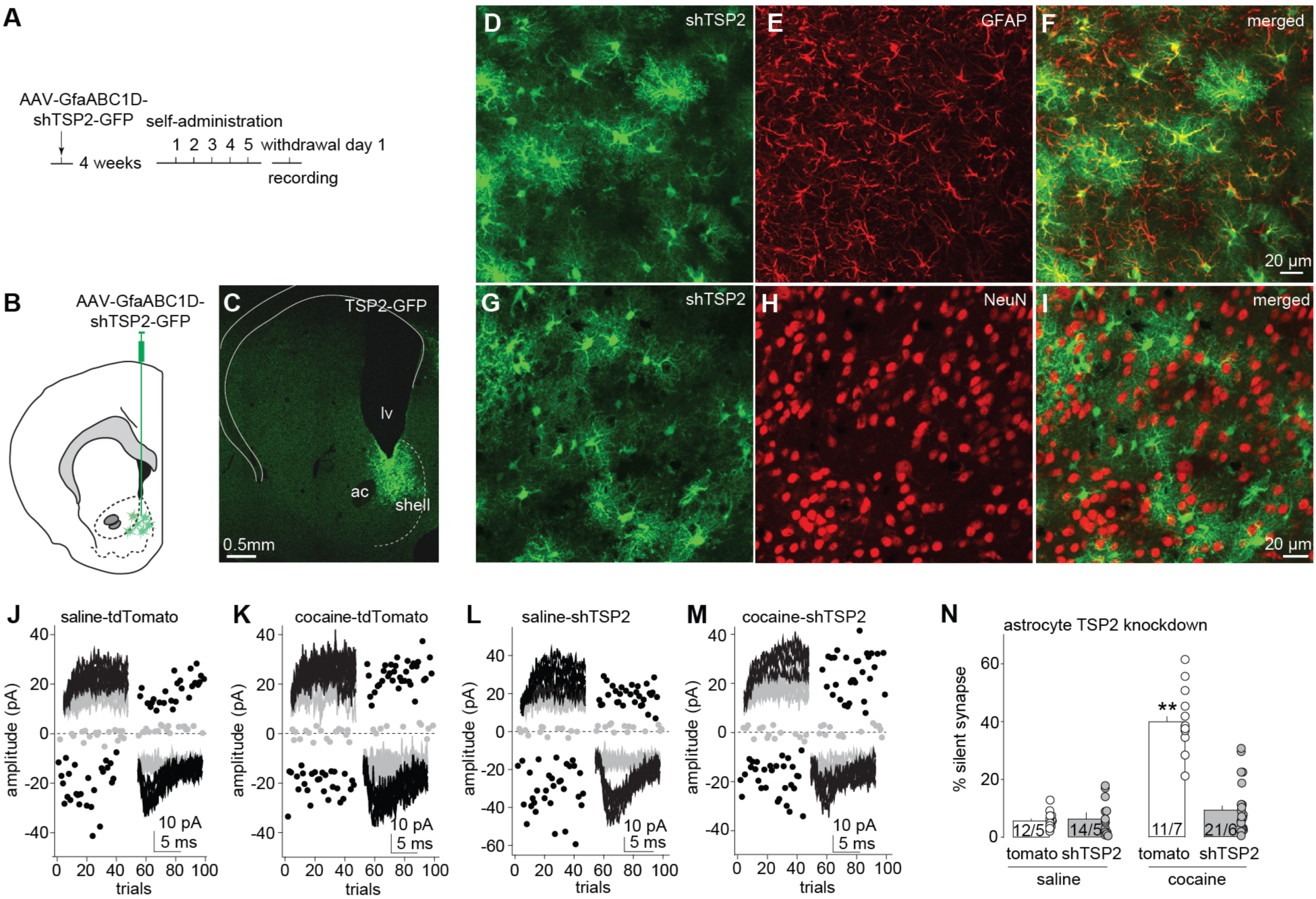
Cocaine-induced generation of silent synapses requires astrocytic TSP 2. **A** Scheme of experimental procedure illustrating that rats first received intra-NAcSh injection of AAV that expressed shTSP2 under the GfaABC_1_D promotor, then received 5-day saline or cocaine self-administration training, and were sacrificed for silent synapse analysis 1 day after the training procedure. **B,C** Diagram (**B**) and confocal image showing the GfaABC_1_D-mediated expression (**C**) of shTSP2 in the NAcSh. **D-F** Confocal images of an example NAcSh slice showing expression of shTSP2 (green) within cells with bushy, astrocyte-like morphologies (**D**), and GFAP staining (red) of the same slice for astrocytic skeletons (**E**) that were overlapped with green signals (**F**). G-I Confocal images of an example NAcSh slice showing expression of shTSP2 (green; **G**), and co-staining with NeuN (red; **H**) revealing minimal overlap between shTSP2-expressing cells and neurons (**I**). **J-M** Example trials of the minimal stimulation assays of NAcSh MSNs from saline (**J**)- or cocaine-trained (**K**) rats with NAcSh astrocytic expression of tdTomato alone, or saline (**L**) or cocaine-trained (**M**) rats with NAcSh astrocytic expression of shTSP2-GFP. Insets showing example EPSCs at +40 or −70 mV from the minimal stimulation assay. **N** Summary showing that astrocytic knockdown of TSP2 did not affect the basal % silent synapses in saline-trained rats, but was sufficient to prevent generation of silent synapses in cocaine-trained rats (F_1,19_ = 91.07, p < 0.01, animal-based two-way ANOVA; p < 0.01 saline-tdTomato vs. cocaine-tdTomato, p = 1 saline-tdTomato vs. saline-shTSP2, p<0.01 cocaine-tdTomato vs. cocaine-shTSP2, p = 1 saline-shTPS2 vs. cocaine-shTSP2, Bonferroni posttest). **, p < 0.01.

We then trained rats to self-administer saline or cocaine 4 weeks after they received bilateral intra-NAcSh injection of shTSP2-GFP-expressing AAV, using rats with AAV expression of tdTomato alone as controls (**Fig. S3**). On withdrawal day 1, we performed the minimal stimulation assay in NAcSh MSNs within the fluorescence-indicated astrocytic matrix. In tdTomato rats, cocaine-induced generation of silent synapses in NAcSh MSNs remained intact, indicating that the AAV transduction per se did not influence the effect of cocaine (**Fig. 3J,K,N**). Furthermore, NAcSh MSNs exhibited a similarly low % silent synapses in saline-trained rats with either astrocytic expression of shTSP2 or tdTomato alone, indicating that knocking down TSP2 did not affect the basal % silent synapses (**Fig. 3J,L,N**). However, in cocaine-trained rats with astrocytic expression of shTSP2, the % silent synapses of NAcSh MSNs was not increased, but comparable to saline-trained rats (**Fig. 3M,N**). These observations show that astrocyte-derived TSP2 is essential for cocaine-induced generation of silent synapses.

### Role of α2δ-1

α2δ-1, known to mediate the actions of gabapentin (GBP) and related drugs, serves as the canonical neuronal signaling substrate for TSP-mediated synaptogenesis during development (Allen and Eroglu, 2017; Eroglu et al., 2009; Risher et al., 2018). We therefore explored whether α2δ-1 also served as an essential signaling component for cocaine-induced generation of silent synapses using the 5-day repeated i.p. injection procedure. Around 1 h before cocaine injection each day, mice received i.p. injection of GBP at a dose (80 mg/kg) that effectively inhibits α2δ-1 and prevents TSP-α2δ-1-mediated synaptogenesis in vivo (Eroglu et al., 2009; Risher et al., 2018). Three other groups of mice were used as controls, including rats with repeated i.p. injections of vehicle before saline, GBP before saline, and vehicle before cocaine (**Fig. 4A**). One day after the drug procedure, cocaine-treated mice without GBP administration exhibited increased % silent synapses in NAcSh MSNs compared to saline-treated mice (**Fig. 4B-F**). This increase of % silent synapses by cocaine was prevented by GBP pre-injection (**Fig. 4B-F**), implicating α2δ-1 in cocaine-induced generation of silent synapses.

**Figure 4.**
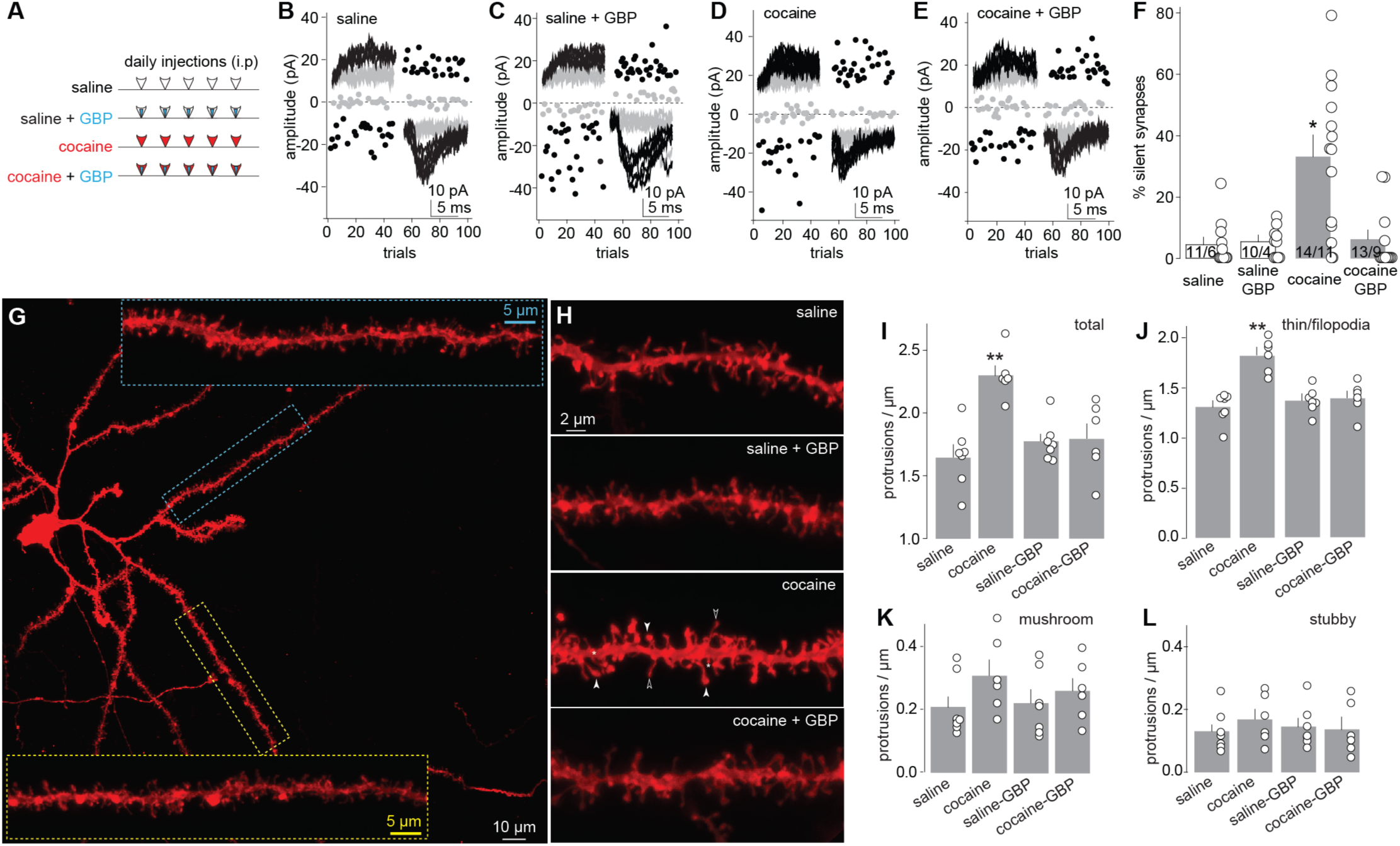
Inhibition of α2δ-1 by GBP prevents cocaine-induced generation of silent synapses and spinogenesis. **A** Experimental schemes showing a 2-by-2 design (repeated i.p. injections of saline or cocaine versus co-administration of vehicle or GBP) in 4 animal groups. **B-E** Example trials of the minimal stimulation assays of NAcSh MSNs from saline-exposed mice with co-administration of vehicle (**B**) or GBP (**C**) mice, and from cocaine-exposed mice with co-administration of vehicle (**D**) or GBP (**E**). Insets showing example EPSCs at +40 or −70 mV from the minimal stimulation assay. **F** Summary showing that co-administration of GBP did not affect the basal % silent synapses in saline-exposed mice, but prevented generation of silent synapses in cocaine-exposed mice (F_1,26_ = 5.69, p=0.03, animal-based two-way ANOVA; p < 0.01 saline vs. cocaine, p = 1.00 saline vs. cocaine-GBP, Bonferroni posttest). **G**-**H** Confocal images of Di-based staining of an example NAcSh MSN with extensive dendrites (**G**) and example MSN dendrites from rats with 4 different treatment described in **A** (**H**). **I-L** Summaries showing that one day after the repeated i.p. cocaine procedure, the densities of total spines (F_1,22_ = 14.17, p <0.01, animal-based two-way ANOVA; p < 0.01 saline vs. cocaine, p = 1.00 saline vs. cocaine-GBP, Bonferroni posttest; **I**) and thin-filopodia-like spines (F_1,22_ = 11.5, p < 0.01, animal-based two-way ANOVA; p < 0.01 saline vs. cocaine, p = 1.00 saline vs. cocaine-GBP; **J**), but not mushroom-like (F_1,22_ = 1.11, p = 0.30, animal-based two-way ANOVA; **K**) or subby spines (F_1,22_ = 1.07, p = 0.31, animal-based two-way ANOVA; **L**), were increased in NAcSh MSNs, and these effects of cocaine were prevented by co-administration of GBP. * P < 0.05; ** P < 0.01.

A prominent prior result aligning cocaine-generated silent synapses to new synapses is the concurrent increase in the density of relatively immature, thin spines on the dendrites of NAcSh MSNs (Brown et al., 2011; Graziane et al., 2016; Wright et al., 2019). We therefore used the GBP co-administration strategy to examine whether inhibiting TSP2-α2δ-1 signaling prevented cocaine-induced spinogenesis. As established previously (Brown et al., 2011; Graziane et al., 2016), we used DiI-based imaging to sample secondary dendrites where NAcSh MSNs form dense spines for receipt of excitatory synaptic projections (**Fig. 4G**). One day after the self-administration training, cocaine-treated rats without co-administration of GBP exhibited higher densities of total dendritic spines compared to saline-trained rats (**Fig. 4H,I**). This increase was primarily attributable to increases in the density of thin/filopodia-like spines, and not mushroom-like or stubby spines (**Figs. 4J-L; S4**). This result confirms the correlation between cocaine-generated silent synapses and thin/filopodia-like spines under our current experimental conditions. GBP co-administration prevented cocaine-induced increases in the density of thin/filopodia-like spines (**Fig. 4H-L**). These findings demonstrate that an α2δ-1-mediated synaptogenic mechanism is essential for both cocaine-induced generation of silent synapses and increases in spine densities, thus, mechanistically linking cocaine-generated synapses to newly generated spines in the NAcSh.

### Role of postsynaptic α2δ-1

α2δ-1 is expressed both pre- and postsynaptically. To determine where the synaptogenic effects of α2δ-1 take place, we designed and packaged an shRNA (shα2δ-1) together with a GFP expression cassette into AAV2/9 (**Fig. S5A,B**). We first performed postsynaptic manipulations by bilaterally injecting the rats with shα2δ-1-GFP-expresing AAV into the NAcSh, allowing knockdown of α2δ-1 in postsynaptic MSNs. Rats with intra-NAcSh injection of an AAV2/9 that expressed GFP alone were used as controls. Four weeks later, we trained the rats for saline or cocaine self-administration (**Figs. 5A; S5C-E**). On withdrawal day 1, we prepared the NAcSh slices and analyzed silent synapses in GFP-positive NAcSh MSNs by stimulating presumably non-transduced excitatory presynaptic fibers (**Fig. 5B-D**).

**Figure 5.**
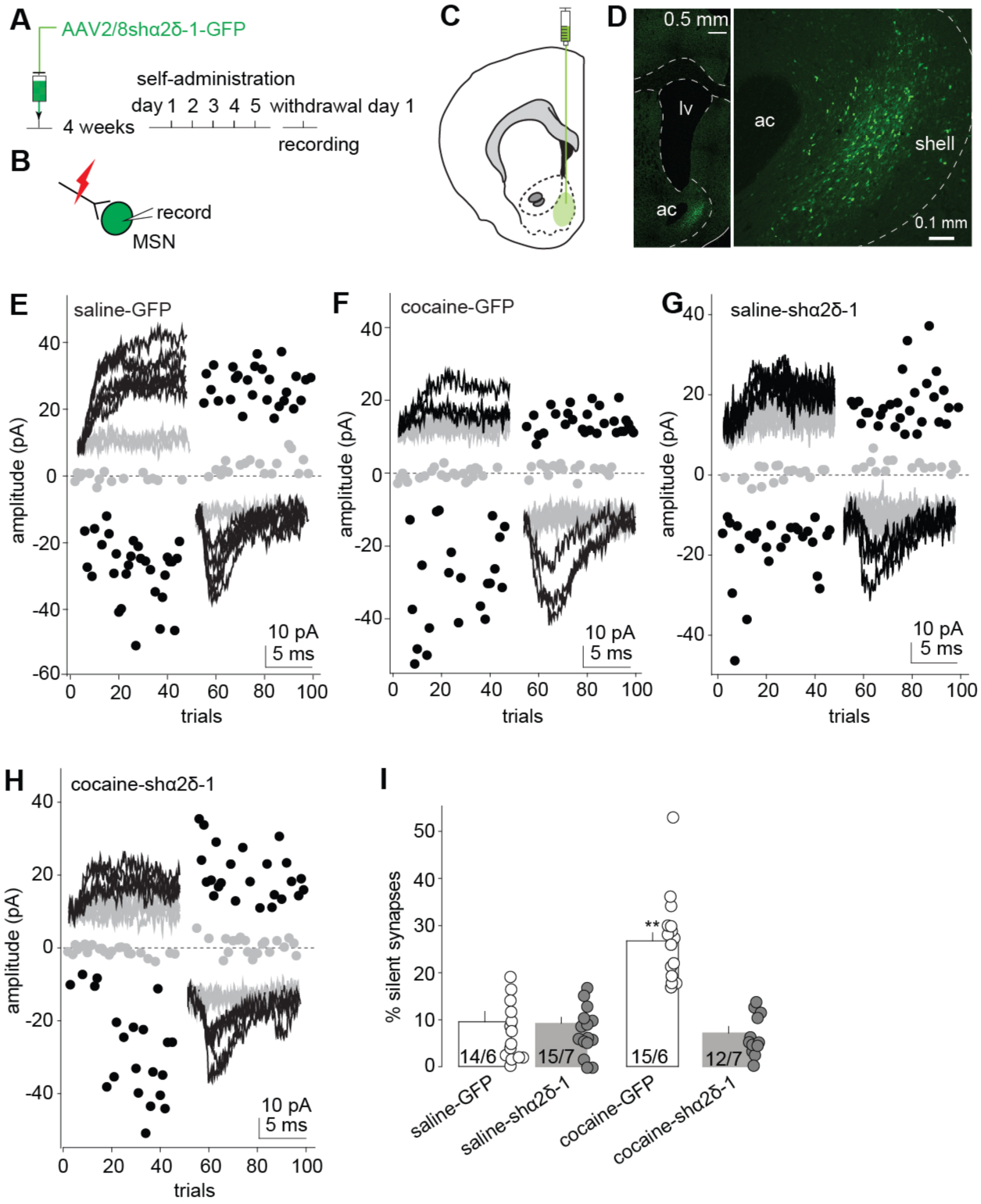
Knockdown of postsynaptic α2δ-1 prevents cocaine-induced generation of silent synapses. **A** Experimental schemes showing that four weeks after receiving intra-NAc injection of shα2δ-1-expressing AAV, rats were trained for saline or cocaine self-administration, followed by the minimal stimulation assay on withdrawal day 1. **B** Diagram showing that virally infected MSNs, identified by their GFP signals, were selectively recorded. **C,D** Diagram (**C**) and example confocal images (**D**) showing the injection site and viral expression. **E-H** Example trials of the minimal stimulation assays of NAcSh MSNs that expressed GFP alone from saline (**E**)- or cocaine-trained (**F**) rats, or MSNs that expressed shα2δ-1-GFP from saline (**G**)- or cocaine-trained (**H**) rats. Insets showing example EPSCs at +40 or −70 mV from the minimal stimulation assay. **I** Summary showing that postsynaptic expression of shα2δ-1 did not affect the basal % silent synapses in saline-trained rats, but prevented generation of silent synapses in cocaine-trained rats (F_1,22_ = 34.30, p < 0.01, animal based two-way ANOVA; p < 0.01 saline-GFP vs. cocaine-GFP, p = 1 saline-GFP vs. saline-shα2δ-1, p < 0.01 cocaine-GFP vs. cocaine-shα2δ-1, p = 1 saline-shα2δ-1 vs. cocaine-shα2δ-1, Bonferroni posttest). ** P < 0.01.

In saline-trained rats, NAcSh MSNs expressing GFP or shα2δ-1-GFP exhibited similarly low % silent synapses, indicating that viral expression of shα2δ-1 per se did not affect the basal % silent synapses (**Fig. 5E-I**). However, in cocaine-trained rats, while NAcSh MSNs that expressed GFP alone exhibited a high % silent synapses as expected, MSNs that expressed shα2δ-1 exhibited a low % silent synapses at levels comparable to saline-trained rats (**Fig. 5E-I**). These results indicate that postsynaptic α2δ-1 is essential for the generation of silent synapses in response to cocaine.

### Role of presynaptic α2δ-1

To examine presynaptic α2δ-1, we first established a presynaptically specific manipulation. We focused on synapses within the projection from the infralimbic prefrontal cortex (IL) to NAcSh, in which silent synapses are generated by cocaine self-administration (Ma et al., 2014). Our strategy was to co-express channelrhodopsin2 (ChR2) with shα2δ-1 in IL neurons, such that EPSCs triggered by optogenetic stimulation in NAcSh slices were evoked from the IL presynaptic fibers with α2δ-1 knockdown (**Fig. 6A**). To achieve the co-transduction, we employed a double-virus system, consisting of an AAV2/8 that expressed ChR2-mCherry and another AAV2/8 that expressed shα2δ-1-GFP. We mixed the two viruses at a 1:1 ratio for co-injection, a strategy that achieves sufficient co-transduction in our previous electrophysiological and behavioral studies (Neumann et al., 2016; Wang, 2020). We bilaterally injected the rats with this viral mixture into the IL (**Fig. 6A,B**). Four weeks later, mCherry and GFP signals were enriched within the IL (**Fig. 6C-E**).

**Figure 6.**
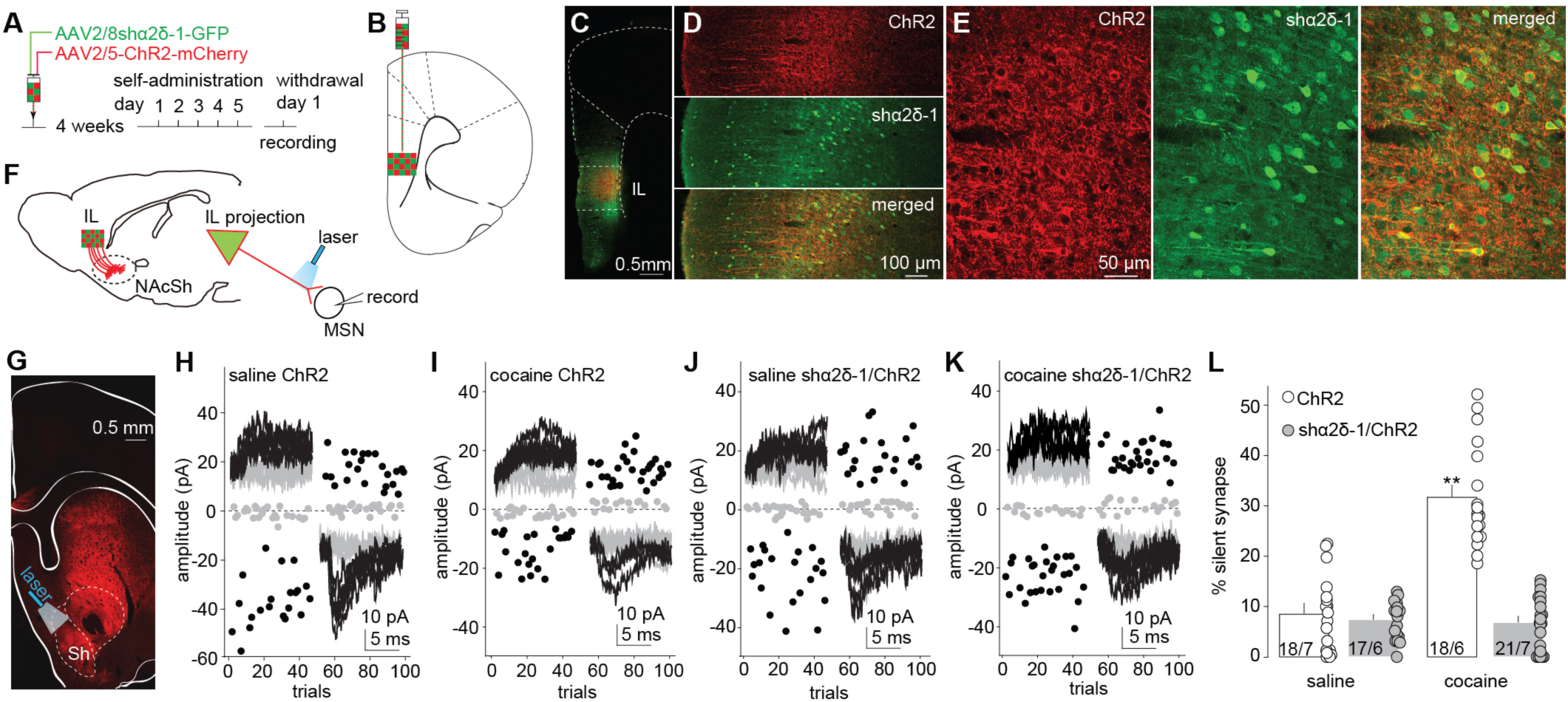
Knockdown of presynaptic α2δ-1 prevents cocaine-induced generation of silent synapses. **A** Experimental schemes showing that rats were co-injected with the shα2δ-1-expressing and ChR2-expressing AAVs in the mPFC, then trained for saline or cocaine self-administration, followed by the minimal stimulation assay on withdrawal day 1. **B,C** Diagram (**B**) and example confocal images showing the viral injection site infralimbic mPFC (IL) (**C**). **D,E** Confocal images of an example IL slice showing co-expression of shα2δ-1 (green) and ChR2 (red) at low (**D**) and high (**E**) magnifications. **F** Diagram illustrating the optogenetic minimal stimulation assay through which silent synapses were sampled from the IL-to-NAc transmission in which the IL projections co-expressed ChR2 and shα2δ-1. **H-K** Example trials of the optogenetic minimal stimulation assays of NAcSh MSNs from saline (**H**)- or cocaine-trained (**I**) rats with intra-IL expression of ChR2 alone, saline (**J**)- or cocaine-trained (**K**) rats with co-expression of ChR2 and shα2δ-1 in the IL. Insets showing example EPSCs at +40 or −70 mV from the minimal stimulation assay. **L** Summary showing that presynaptic expression of shα2δ-1 did not affect the basal % silent synapses in saline-trained rats, but prevented generation of silent synapses in cocaine-trained rats (F_1,22_ = 44.68, p < 0.01, animal based two-way ANOVA; p < 0.01 saline-ChR2 vs. cocaine-ChR2, p = 1 saline-ChR2 vs saline-ChR2-shα2δ-1, p < 0.01 cocaine-ChR2 vs. cocaine-ChR2-shα2δ-1, p = 1 saline-ChR2-shα2δ-1 vs cocaine-ChR2-shα2δ-1, Bonferroni posttest).

We trained rats with intra-IL expression of ChR2-mCherry and shα2δ-1-GFP to self-administer saline or cocaine, using the rats with intra-IL expression of ChR2-mCherry alone as controls (**Figs. 6A; S6A-C**). One day later, we performed the optogenetics-based minimal stimulation assay, as previously demonstrated (Lee et al., 2013; Ma et al., 2014), to selectively examine the % silent synapses within the IL-to-NAcSh projection (**Figs. 6G; S6D-F**). In saline-trained rats, a similarly low % silent synapses was observed within the IL-to-NAcSh projections with or without presynaptic expression of shα2δ-1, suggesting that presynaptic knockdown of α2δ-1 did not, by itself, affect basal levels of silent synapses (**Fig. 6H-L**). In cocaine-trained rats, the IL-to-NAcSh projection that only expressed ChR2 exhibited an increased % silent synapses as expected; however, this increase was prevented by presynaptic co-expression of shα2δ-1 (**Fig. 6H-L**). As an interleaved control, we also performed electrical stimulation, which extensively included presynaptic terminals that did not express shα2δ-1 andChR2 (**Fig. S6E**). Under this condition, we observed an increased % silent synapses in cocaine trained rats (**Fig. S6F,G**). Thus, presynaptic α2δ-1 is also essential for cocaine-induced generation of silent synapses. Such an essential role of presynaptic α2δ-1 may be unique for cocaine-induced synaptogenesis in the adult NAcSh, as in other brain regions during development presynaptic α2δ-1 is not involved in TSP2-induced synaptogenesis (Risher et al., 2018) (see Discussion).

### Behavioral correlates

The above mechanistic link between TSP-α2δ-1 signaling and cocaine-induced generation of silent synapses argues that these silent synapses are nascent, immature synaptic contacts generated de novo to establish new circuit connections in the NAcSh that encode memory traces associated with cocaine experience. If so, their presence may promote cocaine seeking after withdrawal. To test this, we focused on cue-induced cocaine seeking, which results from cue-conditioned cocaine self-administration training and intensifies after cocaine withdrawal (Grimm et al., 2001).

We first took a focal approach to selectively disrupt TSP-α2δ-1 signaling in the NAcSh. Specifically, we bilaterally injected rats with the shα2δ-1-GFP-expressing AAV into the NAcSh, using rats with intra-NAcSh injection of GFP-expressing AAV as controls (**Fig. 7A**). Four weeks later, these rats underwent cocaine self-administration, during which both the shα2δ-1 and GFP rats established cue-conditioned nose poke responding for cocaine infusions (**Figs. 7B,C; S7A-H**). After 45-d withdrawal, the rats were placed in a 1-h cocaine seeking test, in which nose pokes resulted in the presence of light cues but not cocaine infusion. The number of nose pokes serves as a measure for cue-induced cocaine seeking, driven by cue-associated cocaine memories. Compared to control GFP rats, shα2δ-1 rats exhibited reduced cue-induced cocaine seeking (**Figs. 7D; S7A-H**).

**Figure 7.**
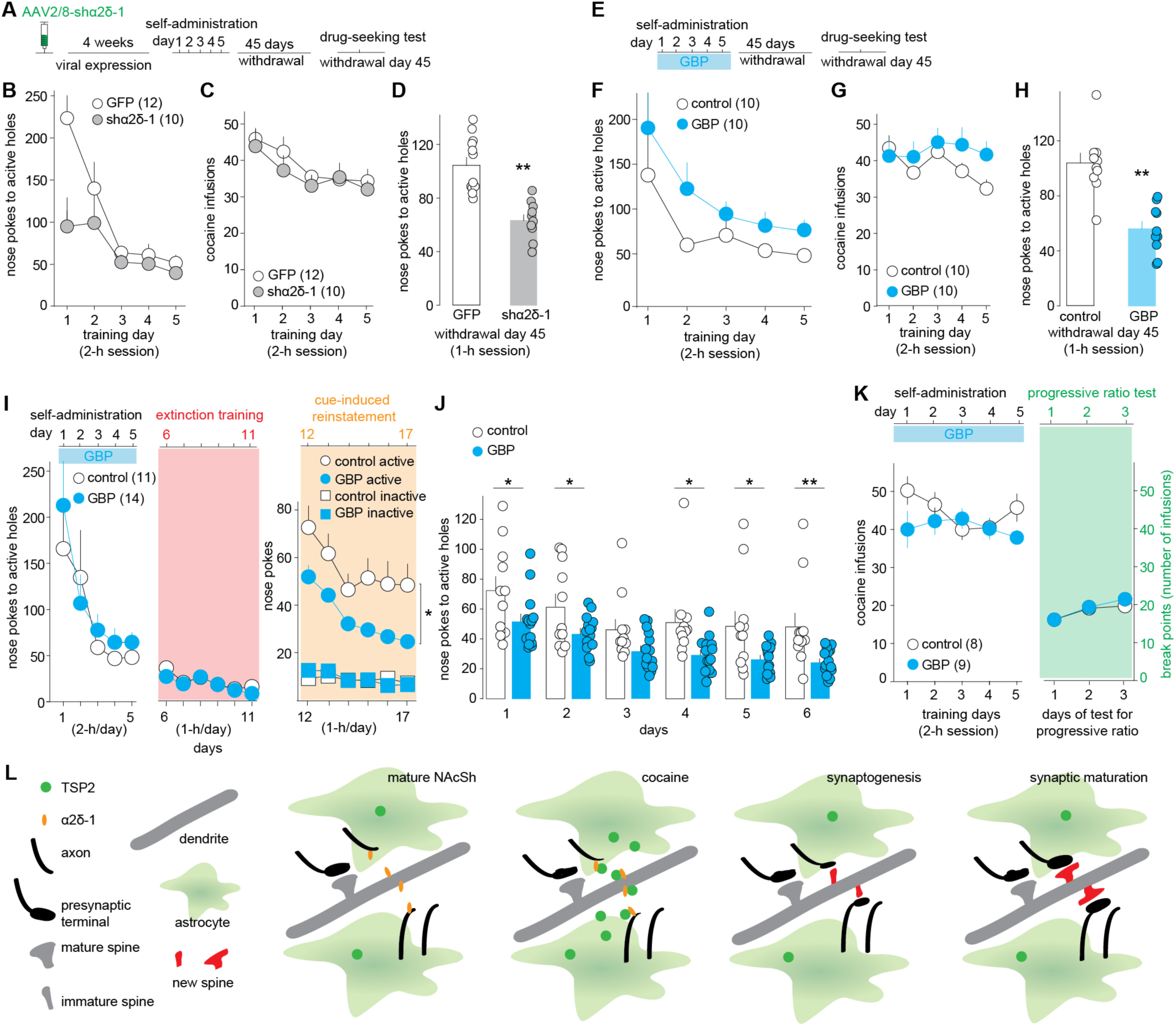
NAcSh synaptogenesis contributes to cue-associated cocaine memories. **A** Experimental schemes illustrating that rats with intra-NAcSh expression of shα2δ-1 were trained for cocaine self-administration, followed by a cue-induced cocaine seeking test on withdrawal day 45. **B,C** Summarized results showing nose poke responding (**B**) and cocaine infusion (**C**) of rats with Intra-NAcSh expression of GFP or shα2δ-1-GFP over the 5-day cocaine self-administration procedure. **D** Summary showing that, compared to rats with intra-NAcSh expression of GFP, rats with intra-NAcSh expression of shα2δ-1-GFP exhibited lower levels of cue-induced nose poke responding during the 1-h extinction test on withdrawal day 45 (t_1,20_ = 5.37, p < 0.01, t-test). **E** Experimental schemes illustrating that rats are co-administrated with either vehicle or GBP during the 5-day cocaine self-administration procedure, followed by a cue-induced cocaine seeking test on withdrawal day 45. **F,G** Summary of nose poke responding (**F**) and cocaine infusion (**G**) of rats with co-administration of vehicle or GBP over the 5-day cocaine self-administration procedure. **H** Summary showing that, compared to cocaine-trained rats with vehicle co-administration, cocaine-trained GBP rats exhibited lower levels of cue-induced nose poke responding during the 1-h cue-induced cocaine seeking test on withdrawal day 45 (t_1,18_ = 5.28, p < 0.01, t-test). **I** Summaries showing that, compared to vehicle rats, GBP rats exhibited similar levels of nose poke responding during the 5-day cocaine self-administration procedure (left), similar rates of extinction of operant responding (F_1, 23_ = 3.30, p = 0.08, between subject effect of GBP, mixed ANOVA with repeated measures; middle), but decreased levels of cue-induced reinstatement of cocaine seeking (right) (F_1, 23_ = 7.84, p = 0.01, effect of GBP, mixed ANOVA with repeated measures). **J** Summary showing that, compared to rats with co-administration of vehicle during cocaine self-administration, rats with co-administration of GBP exhibited reduced nose poke responding over the 6-day tests of cue-induced reinstatement of cocaine seeking (day 1, t_1,23_ = 2.11, p < 0.05; day 2, t_1,23_ = 2.13, p < 0.05; day 3, t_1,23_ = 2.02, p = 0.06; day 4, t_1,23_ = 2.66, p < 0.05; day 5, t_1,23_ = 2.46, p < 0.05; day 6, t_1,23_ = 2.87, p < 0.01, two-tailed t-test). **K** Summary showing that rats with co-administration of vehicle or GBP exhibited similar levels of cocaine infusion during cocaine self-administration and breaking points after self-administration training (F_1,15_ = 0.30, p = 0.59, effect of GBP, mixed ANOVA with repeated measures). **L** Diagrams summarizing our results that cocaine experience induces astrocytic release of TSP2 and activation of both pre- and postsynaptic α2δ-1, resulting in generation of new, silent synapses, some of which fully mature after cocaine withdrawal.

While the above AAV-mediated manipulations are specific to NAcSh neurons, it was unclear whether such a long-term knockdown of α2δ-1 affected memory formation during the self-administration training or expression of the memories during the cocaine seeking test. To address this, we took a pharmacological approach and treated rats with daily GBP injections during the 5-day cocaine self-administration period, an approach that prevented TSP-α2δ-mediated generation of NAcSh silent synapses after cocaine (**Fig. 4**). The rats with daily injections of saline vehicle were used as controls. Despite increased variability in acquisition curves, which might be associated with the global effect of GBP, both GBP and vehicle rats established cue-conditioned nose poke responding, resulting in similar levels of cocaine consumption (**Figs. 7E-G; S7I-N**). We then placed the rats back in their home cages for drug withdrawal without additional GBP administration. During the 1-h cocaine seeking test on withdrawal day 45, GBP rats exhibited substantially decreased levels of cue-induced cocaine seeking compared to vehicle rats (**Fig. 7H**). These results specifically implicate TSP-α2δ-1-mediated NAc synaptogenesis during cocaine self-administration as a key mechanism contributing to the formation of memory traces that drive cue-induced cocaine seeking.

While cocaine seeking is a complex and multifaceted behavior, the above results highlight the cue-associated component. To further focus on discrete cues, we employed a cue-induced reinstatement of cocaine seeking test. Through cue-conditioned cocaine self-administration training, both GBP and vehicle rats established nose poke responding for cocaine infusion in a similar manner (**Fig. 7I, S7O**). This result, together with the results from rats with intra-NAcSh expression of shα2δ-1 (**Fig. 7B,C**), suggests that NAcSh synaptogenesis is not required for acquiring the operant responding of cocaine self-administration. Starting at withdrawal day 1, both groups of rats then underwent a 6-day extinction training (1h/d), during which nose-poking to cocaine-predicting holes (i.e., active holes) resulted in neither cocaine infusions nor light cues. As such, operant responding was preferentially extinguished while the cue-cocaine association was not targeted, and thus was thought to remain largely intact. Both GBP and vehicle rats exhibited similar extinction rates, suggesting minimal involvement of NAcSh synaptogenesis in operant responding per se (**Fig. 7I**). The rats were then placed into the same operant boxes for a 6-d test (1h/d) of cue-induced reinstatement of cocaine seeking, during which nose-poking to the cocaine-predicting hole still did not lead to cocaine infusion but resulted in presentation of the light cue, thus setting the cue-cocaine association as a primary driving factor for the operant responding. Over the 6 days of testing, GBP rats exhibited consistently lower levels of operant responding compared to vehicle rats (**Figs. 7I,J; S7O,P**), further suggesting that NAcSh synaptogenesis and silent synapse generation likely underlie the cue-associated cocaine memories (see Discussion).

In contrast, we did not detect changes in the motivational aspect of cocaine seeking upon disruption of NAcSh synaptogenesis. Specifically, one day after training GBP- and vehicle-treated rats with 5-d cocaine self-administration on the fixed ratio 1 schedule, we placed the rats back in the operant boxes for cocaine self-administration on a progressive ratio schedule (see **Methods**), in which a progressively increased number of nose pokes was needed to obtain the next cocaine infusion. The highest number of cocaine infusions that the rats obtained before giving up, referred to as the break point, reflects the level of motivation to obtain cocaine (Gardner, 2000; Richardson and Roberts, 1996). In all 3 consecutive progressive ratio tests, GBP and vehicle rats exhibited similar break points, a result that does not implicate NAcSh synaptogenesis in forming the motivational representation of cocaine during early withdrawal periods (**Figs. 7K; S7Q**).

Collectively, while preventing synaptogenesis-mediated generation of silent synapses did not affect the acquisition of cocaine self-administration per se, it compromised cue-induced cocaine seeking after drug withdrawal and cue-induced reinstatement of cocaine seeking after extinction. Thus, rather than the entire cocaine-associated memory, NAcSh silent synapses are involved selectively in specific components of the cocaine memories.

## Discussion

During brain development, astrocytic TSP-α2δ-1 signaling serves as a key mechanism promoting synaptogenesis (Allen and Eroglu, 2017). Our current results suggest that this developmental mechanism is employed by cocaine experience to generate nascent, AMPAR-silent synapses in the adult NAcSh, with the resulting synaptic and circuit remodeling promoting cue-induced cocaine seeking and relapse.

### Cocaine-induced synaptogenesis

Prior to this current study, two lines of evidence suggest that exposure to cocaine generates new excitatory synapses in the NAc. First, after repeated exposure to cocaine, the density of dendritic spines, the main synaptic targets of excitatory axonal terminals (Yuste and Denk, 1995), is increased in NAc MSNs (Robinson and Kolb, 1999), suggesting spinogenesis and synaptogenesis. However, dendritic spines in vivo exhibit high turnover rates in certain brain regions (Attardo et al., 2015). Thus, the increased spine density after cocaine may simply reflect a shift in the balance between spine generation and elimination. Nonetheless, time-lapse two-photon imaging detects spine generation in NAc slices in response to co-application of dopamine D1 receptor agonists and glutamate, a pharmacological proxy for cocaine’s effects (Dos Santos et al., 2017), potentially linking spinogenesis to cocaine-induced increases in NAc spine density. Second, repeated exposure to cocaine increases the level of AMPAR-silent excitatory synapses in the NAc (Huang et al., 2009). These synapses share key features with immature, silent synapses that are abundant in the developing brain (Hanse et al., 2013; Huang et al., 2015a; Kerchner and Nicoll, 2008). Furthermore, cocaine-induced generation of silent synapses is achieved by a series of pro-synaptogenic processes, such as synaptic expression of GluN2B-containing NMDARs and activation of CREB (Brown et al., 2011; Huang et al., 2009; Marie et al., 2005; Russo et al., 2010), which, among other factors, are critically implicated in synaptogenesis and circuit formation during brain development (Cull-Candy and Leszkiewicz, 2004; Lonze and Ginty, 2002; Shipton and Paulsen, 2014). These results lead to the hypothesis that cocaine-generated silent synapses are nascent synapses that function to rewire NAc circuits (Dong and Nestler, 2014; Russo et al., 2010). Our current study mechanistically links these two lines of results together. As hypothesized, exposure to cocaine activates astrocytic synaptogenic processes by releasing TSP2, which, in turn, activates its neuronal receptor, α2δ-1, to generate new excitatory synapses in the NAc. Disrupting this synaptogenic signaling pathway by inhibiting α2δ-1 simultaneously prevented the cocaine-induced increase in the % silent synapses and the density of dendritic spines (**Fig. 4**). Thus, generated through shared mechanisms, silent synapses and increased spines likely represent the same set of synapses detected through functional versus anatomical approaches.

Although a defining biomarker of cocaine-generated synapses has not been identified, these synapses possess traits that distinguish them from synapses pre-existing before cocaine experience. A signature trait is the dynamics of GluN2B-containing NMDARs, which are enriched when these synapses are generated initially (Brown et al., 2011; Huang et al., 2009), but are replaced by nonGluN2B-, likely GluN2A-, containing NMDARs upon maturation after long-term withdrawal from cocaine (Wright et al., 2019). This compositional switch of NMDARs echoes the maturation of excitatory synapses during development, during which this switch allows AMPARs to be inserted and stabilized (Adesnik et al., 2008; Cull-Candy and Leszkiewicz, 2004; Quinlan et al., 1999). Another cellular trait is the dynamics of calcium-permeable AMPARs (CP-AMPARs). Upon maturation, cocaine-generated silent synapses recruit CP-AMPARs, which are only sparsely expressed at excitatory synapses on NAc MSNs in drug-naïve animals (Lee et al., 2013; Ma et al., 2014; Wright et al., 2019). However, CP-AMPARs are internalized from these synapses when cocaine-associated memories are reactivated, and are re-inserted to these synapses upon reconsolidation of cocaine memories (Wright et al., 2019). Thus, despite maturation after cocaine withdrawal, AMPARs at these synapses remain labile to a certain extent and are sensitive to further manipulations. This implies another potential cellular trait of these synapses, namely, low levels of PSD95 and other synaptic scaffolds. PSD95 functions to stabilize AMPARs at mature synapses, while genetic deletion or knockdown of PSD95 increases the level of AMPAR-silent synapses, potentially resulting from increased lability of AMPARs (Beique et al., 2006; Favaro et al., 2018; Shukla et al., 2017). At the signaling level, TSP-α2δ-1-mediated synaptogenesis in developing cortical neurons requires the small GTPase Rac1 (Risher et al., 2018). After withdrawal from cocaine, neuronal levels of active Rac1 control the transition between the silent versus mature states of cocaine-generated synapses during cocaine memory destabilization and reconsolidation (Wright et al., 2019). Thus, it is possible that the levels or subcellular location of active Rac1 are pre-tuned during synaptogenesis to act as the key signaling molecule that dictates the functional states of cocaine-generated synapses thereafter.

### Astrocytic signaling

In the developing brain, formation and maturation of new synapses critically rely on astrocyte-derived signals (Allen and Eroglu, 2017). Among these, astrocyte-secreted TSP1 and 2, two TSP isoforms that share the same functional domains, contribute to the extracellular matrix around neuronal processes (Adams, 2001; Bornstein et al., 2004), and induce formation of AMPAR-silent glutamatergic synapses (Christopherson et al., 2005; Eroglu et al., 2009). After development, astrocytic Ca^2+^ activities and TSP expression are greatly downregulated (Allen and Eroglu, 2017; Iruela-Arispe et al., 1993; Kim et al., 2017). However, strong astrocytic Ca^2+^ oscillations and increased TSP1/2 expression are observed in the adult cortex after brain damage (Ding et al., 2007; Kuchibhotla et al., 2009; Lin et al., 2003; Nagai et al., 2019), suggesting that TSP mechanisms are merely dormant and capable of being activated to reinitiate synaptogenic processes in the adult brain. Indeed, we show that, after cocaine experience, synaptogenic TSP-α2δ-1 signaling is employed to generate nascent silent synapses in the NAcSh (**Fig. 7L**). These results are in line with the “neural rejuvenation hypothesis,” which proposes that drug experience reopens certain dormant, developmental mechanisms in the adult brain to restructure and thereby profoundly change key brain circuits involved in addictive behaviors (Dong and Nestler, 2014).

Although low in adult brains, TSP1/2 levels are substantially increased after relatively severe brain insults, such as ischemia- or stroke-induced neurodegeneration, after which massive homeostatic synaptic remodeling is triggered (Liauw et al., 2008; Lin et al., 2003). It will be important to determine whether periods of cocaine administration also change TSP1/2 levels in the adult NAcSh, an experiment that unfortunately requires suitable high-quality antibodies not yet available. α2δ-1 is abundantly expressed in the NAc as well as the cortical regions that project to the NAc in adult rodents (Cole et al., 2005). Thus, should TSP2 be mobilized in sufficient amounts following activation of astrocytes, it could readily stimulate α2δ-1 signaling in the NAc of cocaine-exposed animals.

Among the 5 TSPs, TSP1 and TSP2 share the same trimeric oligomerization and similar domain properties (Adams, 2001; Iruela-Arispe et al., 1993; Lawler et al., 1993; Risher and Eroglu, 2012). TSP1 and TSP2 are simultaneously expressed at peak levels at the start of the synaptogenic period during development, and either of them is sufficient to promote synaptogenesis in astrocyte-free neuronal cultures (Christopherson et al., 2005; Eroglu et al., 2009). Furthermore, while normal synapse numbers are observed in TSP1- or TSP2-deficient mice, synapse density is substantially decreased in TSP1/2 double null mice (Christopherson et al., 2005). Despite the seemingly redundant roles of TSP1/2, studies focusing separately on TSP1 versus TSP2 reveal their potentially differential influence on synaptogenesis. For example, TSP1 expedites the synaptogenic process but does not change the final density of synapses in cultured hippocampal neurons, effects that are mediated primarily by its interaction with neuroligin-1 (Xu et al., 2010), thus without directly involving α2δ-1. In contrast, TSP2 induces robust synaptogenesis in cultured cortical neurons, which is prevented by either pharmacological inhibition or genetic deletion of α2δ-1 (Risher et al., 2018). As such, although sharing many common downstream targets, TSP1 and TSP2 may promote synaptogenesis in a complementary rather than a redundant manner, such that TSP2 promotes the initiation of synaptogenic processes through α2δ-1, while TSP1 facilitates the assembly and stabilization of new synaptic contacts through synaptic adhesion molecules once the synaptogenic process is initiated. In our current study, genetic deletion of TSP2, but not TSP1, prevented cocaine-induced generation of silent synapses (**Figs. 2,3**). These results, together with the results from α2δ-1 manipulations (**Figs. 4-6**), point to TSP2-activated α2δ-1 signaling in cocaine-induced generation of silent synapses.

α2δ-1 is an auxiliary subunit of voltage-gated Ca^2+^ channels, but its synaptogenic effect is likely mediated through the interaction of its VWF-A-like domain with the EGF-like domains of TSPs, independent of Ca^2+^ channels (Eroglu et al., 2009). In the developing cortex, overexpression of α2δ-1 in postsynaptic neurons is sufficient to increase synapse densities, while selective knockout of α2δ-1 in postsynaptic, but not presynaptic neurons, prevents TSP2-induced increases in synapse densities (Eroglu et al., 2009; Risher et al., 2018). These results argue that postsynaptically located α2δ-1 is both sufficient and necessary in TSP-signaling-mediated synaptogenesis. While confirming the essential role of postsynaptic α2δ-1 (**Fig. 5**), our current results demonstrate that presynaptic α2δ-1 is also required for cocaine-induced generation of silent synapses (**Fig. 6**). Given the distinct properties of astrocytes in different brain regions and at different developmental stages (Allen and Eroglu, 2017; Bayraktar et al., 2014; Zhang and Barres, 2010), these results may reflect potentially differential roles of TSP2-α2δ-1 signaling in the cortex versus NAc as well as its differential involvement in developmental versus cocaine-induced synaptogenesis. Alternatively, the results may reveal a unique role of presynaptic α2δ-1 in cocaine-mediated synaptogenesis by NAcSh neurons. Specifically, generation of functional synapses not only requires the formation of pre- and postsynaptic architectures, but also an assembly of functional machineries that mediate presynaptic release and postsynaptic response. α2δ-1 is a key molecular controller that targets P/Q and N-type Ca^2+^ channels (Cav2.1 and Cav2.2) to the presynaptic active zones, a key step that enables Ca^2+^-dependent vesicular release at presynaptic terminals (Catterall and Few, 2008; Hoppa et al., 2012). As such, with the postsynaptic α2δ-1-signaling intact, TSP2 might still have generated new synaptic contacts upon presynaptic knockdown of α2δ-1 (Risher et al., 2018), but these new synapses might not have been presynaptically functional, so that they could not be electrophysiologically detected in the minimal stimulation assay (**Fig. 6**). Such an outcome would be consistent with both Ca^2+^ channel-dependent and independent functions of α2δ-1 being required by TSPs to generate new, functional synapses.

Beyond generation of silent synapses, the TSP1-α2δ-1 signaling is upregulated in rat NAc core after extinction from cocaine self-administration (Spencer et al., 2014), suggesting involvement of astrocytes in multiple time points of cocaine-induced behaviors. Beyond α2δ-1, astrocytic TSPs also bind to a large number of extracellular matrix proteins and cell surface receptors that regulate synaptic remodeling (Risher and Eroglu, 2012). Beyond TSPs, astrocytes also release other factors, such as BDNF, cholesterol, glypicans, hevin, SPARC, TGF-β, and TNF-α that not only promote formation but also refinement of excitatory synapses (Allen and Eroglu, 2017; Kim et al., 2017). Our demonstration that cocaine experience activates astrocytes and reopens the TSP-α2δ-1-mediated developmental mechanism for new synapse formation in the adult NAc (**Fig. 7L**) establishes astrocytes as key cellular substrates that initiate the potential rejuvenation processes through which NAc circuits are redeveloped and rewired to persistently encode drug memories (Dong and Nestler, 2014).

### Silent synapses in cue-induced cocaine seeking

Even when cocaine self-administration is established through relatively straightforward reinforcement schedules as used in our current study, cocaine seeking can be multifaceted, involving many forms of associative learning that collectively drive a large repertoire of cocaine-oriented conditioned and instrumental responding. It is conceivable that NAcSh synaptogenesis and the generation of silent synapses are selectively involved in some, but not all, aspects of cocaine seeking. The fact that preventing TSP-2δ-1-mediated NAcSh synaptogenesis did not affect the acquisition of cocaine self-administration or break points in a progressive ratio test (**Fig. 7**) is not surprising given that excitotoxic lesion of the NAc does not disrupt the acquisition of cocaine self-administration (Ito et al., 2004), nor the instrumental learning that drives goal-directed action (Balleine and Killcross, 1994). Thus, NAcSh synaptogenesis is not likely a key mechanism supporting unconditioned stimulus-driven instrumental drug taking.

Rather, NAcSh synaptogenesis may preferentially contribute to the multiple conditioned and conditioned-to-instrumental associations that form during cocaine self-administration to promote the reinforcing impact of conditioned stimuli (CS). In other words, cue-induced cocaine seeking as a form of CS-conditioned reinforcement requires that an otherwise neutral cue gains reinforcing power through its association with cocaine. Establishing such conditioned reinforcements is thought to rely on conflating stimulus-reward associative processing in the basolateral amygdala (BLA) with the reinforcement processing in the NAc (Everitt, 2014; Mogenson et al., 1980). Cocaine self-administration generates silent synapses within the BLA-to-NAc projection, which mature after cocaine withdrawal and contribute to the progressive intensification of cue-induced cocaine seeking (Lee et al., 2013). Through these new synaptic connections, new informational flow can be established within the BLA-to-NAc projection as part of mechanisms assigning cocaine-conditioned cues with reinforcing impact. Such cue-induced cocaine seeking can be further enhanced by conditioned-instrumental transfer, autoshaping, and other CS-conditioned reinforcement mechanisms (Cardinal et al., 2003; Everitt, 2014), in which synaptogenesis-mediated remodeling of other NAc afferents may contribute, including afferents from the prefrontal cortex and thalamus (Ma et al., 2014; Neumann et al., 2016).

The cue-oriented role of NAcSh synaptogenesis is further demonstrated by the cue-induced reinstatement of cocaine seeking test, in which GBP-treated rats only exhibited decreases in cue-conditioned responding, but not in acquisition, instrumental extinction, or progressive ratio responding for cocaine (**Fig. 7**). These results are consistent with the above proposed role of NAcSh synaptogenesis in CS-conditioned reinforcement. It is worth noting that NAcSh silent synapses as efficient plasticity substrates can also be involved in other learning processes, such as extinction learning and retrieval (Millan et al., 2011), which we did not explicitly test.

### Concluding remarks

Our current results demonstrate that an astrocyte-mediated synaptogenic mechanism reminiscent of development is employed by cocaine experience to generate nascent, immature synapses in the adult NAcSh, promoting the formation of new behavioral patterns. These results reveal astrocytes as active players promoting drug-induced synaptic and circuit remodeling in the formation of cocaine memories. These results also encourage future studies to explore astrocyte-based manipulations in reducing drug memories and drug relapse.

## Materials and methods

### Subjects

Male Sprague-Dawley rats (Charles River), both male and female C57BL/6J wild type mice, and both male and female mice bred from the Aldh1l1-Cre/ERT2 transgenic mouse line (Jackson Laboratory), the ROSA26-Lck-GCaMP6f-flox mouse line, the TSP1 null mouse line (Jackson Laboratory) and the TSP2 null mouse line (Jackson Laboratory) at the age of 8-16 weeks old before surgery were used in all experiments. Rats and mice were housed on a regular 12 h light/dark cycle (light on at 07:00 AM) with food and water available ad libitum. All rats and mice were used in accordance with protocols approved by the Institutional Care and Use Committees at the University of Pittsburgh and Icahn School of Medicine at Mount Sinai.

### Electron microscopy

Rats were perfused through the aorta with 5-10 ml of heparin-saline 100-1000 U/ml, followed by 50 ml of 3.75% acrolein in 2% paraformaldehyde, followed by 200 ml of 2% paraformaldehyde. The fixative was made in 0.1 M phosphate buffer, pH 7.4 (PB). Blocks of fixed brain containing the NAc were cut on a vibratome at 50 µm, and tissue sections were collected in PB. Sections were treated for 30 min in 1% sodium borohydride in PB and then rinsed extensively in PB. Sections were then transferred to 0.1 M tris-buffered saline, pH 7.6 (TBS) before being incubated for 30 min in a blocking solution containing 1% bovine serum albumin, 3% normal goat serum and 0.04% Triton X-100 in TBS. Sections were then incubated for 12-15 h in primary guinea pig antibodies against the vesicular glutamate transporter type 1 (vGlut1; 1: 10,000 to 1:12,000 in blocking solution). The antiserum from Millipore/Sigma (AB5905) was directed against a 19 amino acid synthetic peptide from the C-terminus of the rat vGlut1 protein, and specificity was demonstrated by Western blot analysis (Melone et al., 2005), and by correspondence of immunolabeling patterns with other vGlut1 antisera (Fremeau et al., 2001; Kaneko et al., 2002; Melone et al., 2005; Sakata-Haga et al., 2001). After rinsing, sections were then incubated for 30 min in biotinylated donkey anti-rabbit IgG (Jackson ImmunoResearch Laboratories, West Grove, PA) at 1:400 in blocking solution. After additional rinses, tissue was incubated for 30 min in 1:100 avidin-biotin peroxidase complex (Vectastain Elite kit; Vector Laboratories, Burlingame, CA) and rinsed again. Bound peroxidase was developed by exposure to 0.022% diaminobenzidine (Sigma) and 0.003% hydrogen peroxide in TBS for approximately 5 min before terminating the reaction by buffer rinses.

Tissue preparation for electron microscopy included lipid fixation with osmium tetroxide at 1-2% in PB for 1 h, followed by rinsing in PB. Sections were then dehydrated through increasing concentrations of ethanol in PB followed by propylene oxide. Sections were then transferred to a 1:1 mixture of propylene oxide and epoxy resin (EMBed 812, Electron Microscopy Sciences) and incubated overnight. The following day, tissue was transferred to straight epoxy resin for 2-3 hours. The sections were then carefully laid onto commercial plastic film, coverslipped with an additional plastic sheet, put under heavy weights to keep the tissue flat, and then placed in a 60°C oven until the resin was cured. The NAcSh was identified in resin-cured brain sections, and a small part of this region was trimmed to a trapezoid shape using razor blades. Ultrathin sections at 60 nm were then cut from the surface of these sections and collected onto 400 copper mesh grids. The grids were stained with uranyl acetate and lead citrate to add contrast to the tissue, and the tissue was examined in an FEI Morgagni transmission electron microscope. Random sampling was used to identify axospinous synapses formed by axon varicosities immunoreactive for vGlut1. These synapses were then further examined for evidence of ensheathment by astrocytes. The latter were identified by thin, irregular contours, relatively electron lucent cytoplasm, and occasional formation of tight junctions with other astrocytes.

### Repeated i.p. injection of cocaine and gabapentin

Before drug administration or molecular manipulation, mice were allowed to acclimate to their home cages for at least 7 days. For cocaine intraperitoneal (i.p.) injections, we used a 5-day cocaine procedure, which was similar to earlier studies (Graziane et al., 2016; Huang et al., 2009). Briefly, once a day for five days, rats (∼32 days old) were taken out of the home cage for an intraperitoneal (i.p.) injection of either (-)cocaine HCl (15 mg/kg in saline) or the same volume of saline, and placed back to the home cage immediately. Gabapentin (PHR1049, Sigma) solution (100 mg/ml) was freshly prepared on the same day of the injection, and was i.p. injected to animals at 100 mg/kg, 1 h before i.p. injection of saline or cocaine, or the self-administration session. For animals undergoing self-administration, one extra injection of gabapentin was given 8-12 h after the first gabapentin injection.

### Intravenous surgery

Rats were anesthetized with a xylazine-ketamine mixture (5-10/50-100 mg/kg, i.p.). A silicone catheter was inserted into the jugular vein and passed subcutaneously to the midscapular region and connected to a vascular access button (Instech Laboratories) subcutaneously implanted under the skin. The rats were then single-housed and given 5-7 days to recover before the training session. Catheters were flushed with sterile saline containing gentamycin (5 mg/ml) and heparin (10 units/ml) every 24 h during the recovery and training periods.

### Apparatus and self-administration procedure

Self-administration training and testing were conducted in Plexiglas operant conditioning chambers (20 x 28 x 20 cm; Med Associates, St. Albans, VT) individually encased within a ventilated, sound-attenuating cabinet. A stimulus light was mounted inside the designated active hole. An infusion pump containing a 12-ml syringe was located outside of the cabinet. Tygon tubing connected to the syringe to a liquid swivel (Instech, Plymouth Meeting, PA) suspended above the operant conditioning chamber. The outlet of the swivel was further connected to the vascular access button on the back of the rat via a quick connecting luer. A nose poke to the active hole resulted in a cocaine infusion (0.75 mg/kg per infusion), accompanied by the switch-on of a light within the hole that served as a conditioned stimulus (CS) and a background light. The CS stayed on for 6 sec, and the background light for 20 sec. To avoid overdose, we set the ceiling number of drug infusions was set at 100 for the 12-h overnight sessions and 65 for the 2-h daily sessions.

### Progressive ratio procedure

Following acquisition, rats underwent self-administration according to a progressive ratio (PR) schedule of cocaine reinforcement (1.5 mg/kg/infusion), 5 h in duration every other day until completing 3 PR sessions, with maintenance sessions (0.75 mg/kg/infusion, fixed ratio of 1, 2 h/session) on alternating days. The progressive response requirement was derived from the formula 5*e^(0.2(n+1)), slightly modified from (Richardson and Roberts, 1996), in which “n” represents the number of infusions received during the session. The last completed infusion number before the rat failed to attain another one within 1 h was defined as the break point.

### Extinction of operant responding and cue-induced reinstatement of cocaine seeking

Extinction of operant responding and cue-induced reinstatement of cocaine seeking were conducted in the same operant chambers as self-administration, but sessions only lasted 1 h. In the extinction test, nose pokes to either hole were recorded but did not lead to any consequences. During the reinstatement test, nose poke into the active hole turned on the CS for 6 sec and background light for 20 sec, but without cocaine infusion. Nose pokes into the inactive hole were recorded but triggered no consequences. The extinction session started on the first day after completion of the 5-day self-administration training, and was repeated on daily for 6 consecutive days. One day after the extinction training, the reinstatement test began and lasted for 6 consecutive days.

### Viral vectors

Recombinant adeno-associated viral vectors (AAV) were used for in vivo viral-mediated gene expression. Briefly, the AAV2 genomic back bone expressing venus tagged with the intended gene was pseudotyped with AAV5, 8, or 9 capsid proteins. HEK293T cells were co-transfected with the plasmids pF6 (adenoviral helper plasmid), pRVI, pH21 and the AAV2 plasmid, using linear polyethylenimine assisted transfection (Suska et al., 2013). Cultures grown in DMEM (Biochrom) with 10% substituted FBS (Biochrom, #S0115) were harvested from 15 by 15 cm dishes after 48 h. AAVs were harvested and purified as described previously (Lee et al., 2013; Suska et al., 2013). The shRNA sequences were embedded in miRNA sequences in the 3’UTP and expressed by an RNA Polymerase 2 promoter. The sequence for TSP2 was bwT2a in an miR30 backbone (Fellmann et al., 2013; Weng et al., 2014); for shα2δ-1, it was in an miR155 backbone blTPd (Fowler et al., 2016) or miR30 backbone buTPa (Fellmann et al., 2013). Using this procedure, we in-house generated the AAV2/8-syn-shα2δ-1-GFP (blTPd) or AAV2/9 (buTPa), AAV2/5-GfaABC_1_D-hPMCA2w/b-mCherry, as well as AAV2/8 and AAV2/9 expressing GFP or tdTomato alone. The AAV2/5-hSyn-hChR2 (H134R)-mCherry was purchased from UNC Vector Center, and the AAV2/5-gfaABC_1_D-shTSP2-GFP was custom-made by the UNC Vector center. AAV2/5-gfaABC_1_D-tdTomato was purchased from AddGene (44332).

### Intracranial Viral Infusion

A stereotaxic microinjection technique was used to deliver viral vectors to specific brain regions. Briefly, rats were anesthetized with 1–3% isoflurane gas. Stainless steel cannulae (31 gauge) were implanted bilaterally into the NAcSh (AP: +1.65 mm; ML: ±0.75 mm; DV: −7.65 mm) or the IL-PFC (AP: +3.1 mm; ML: ±0.6 mm; DV: −4.5 mm). Concentrated viral solutions (1 µl/side) were infused through a pump at a flow rate of 0.2 µl /min. To achieve presynaptic knockdown and ChR2 expression in the IL, a mixture of AAV2/8-shα2δ-1-GFP and AAV2/2-hSyn-hChR2(H134R)-mCherry (mixed in a 1:1 ratio, 1 µl/side) was used. The injection cannula was left in place for 5 min and then slowly withdrawn. After the skin was sutured, the rat was placed on a heating pad ∼1 h for postsurgical recovery before being transferred to the home cage.

### Preparation of NAc slices

Rats or mice were decapitated following isoflurane anesthesia. Coronal slices (300 and 350 µm for mice and rats, respectively) containing the NAc were prepared on a VT1200S vibratome (Leica, Germany) in 4°C cutting solution containing (in mM): 135 N-methyl-D-glucamine, 1 KCl, 1.2 KH_2_PO_4_, 0.5 CaCl_2_, 1.5 MgCl_2_, 20 choline-HCO_3_, and 11 glucose, saturated with 95% O_2_ / 5% CO_2_, pH adjusted to 7.4 with HCl. Slices were incubated in artificial cerebrospinal fluid (aCSF) containing (in mM): 119 NaCl, 2.5 KCl, 2.5 CaCl_2_, 1.3 MgCl_2_, 1 NaH_2_PO_4_, 26.2 NaHCO_3_, and 11 glucose, saturated with 95% O_2_ / 5% CO_2_ at 37° C for 30 min and then allowed to recover for >30 min at room temperature before experimentation.

### Electrophysiological recordings

All recordings were made from MSNs located in the NAc shell. During recordings, slices were superfused with aCSF containing 100 µM picrotoxin (P1675, Sigma), which was heated to 31 – 33°C by passing the solution through a feedback-controlled in-line heater (Warner, CT) before entering the chamber. In all EPSC recordings, electrodes (2 – 5 MΩ) were filled with (in mM): 140 CsCH_3_O_3_S, 5 TEA-Cl, 0.4 Cs-EGTA, 20 Hepes, 2.5 Mg-ATP, 0.25 Na-GTP, 1 QX-314, pH 7.3. Picrotoxin (100 µM) was included to inhibit GABAA receptor-mediated currents. For electrically-evoked EPSCs, presynaptic afferents were stimulated by a constant-current isolated stimulator (Digitimer, UK), using a monopolar electrode (glass pipette filled with aCSF). Series resistance was 9 to 20 MΩ, uncompensated, and monitored continuously during recording. Cells with a change in series resistance beyond 15% were not accepted for data analysis. Synaptic currents were recorded with a MultiClamp 700B amplifier, filtered at 2.6–3 kHz, amplified 5 times, and then digitized at 20 kHz. To make recordings of NAcSh MSNs under the influence of virally-infected astrocytes, we chose the recording location with a high density of fluorescent signals suggestive of high densities of infected astrocytes. Furthermore, MSNs that were located within the visible matrix of astrocytic fluorescence were chosen for recordings.

To record EPSCs evoked by optogenetic stimulation, AAV2/5-hSyn-hChR2(H134R)-mCherry (UNC Vector core) was used. For the optogenetic minimal stimulation assay, the laser duration was adjusted between 0.1 - 0.3 ms and the laser power was adjusted at 1-3 mW at the output of the 40x objective lens.

To quantify % silent synapses, we made two theoretical assumptions: 1) the presynaptic release sites are independent, and 2) release probability across all synapses, including silent synapses, is identical. Thus, % silent synapses was calculated using the equation: 1 – ln(F_– 70_)/ln(F_+40_), in which F_–70_ was the failure rate at −70 mV and F_+40_ was the failure rate at +40 mV, as rationalized previously(Liao et al., 1995). Note that in this equation, the failure rate is the only variable that determines the % silent synapses. The amplitudes of EPSCs were used to present failures or successes, but otherwise did not have analytical value. It is possible that aforementioned theoretical assumptions were not always true. Nevertheless, the above equation was still used, as it is valid in predicting the changes of silent synapses qualitatively as previously rationalized (Lee et al., 2013; Ma et al., 2014). The amplitude of an EPSC was determined as the mean value of the EPSC over a 1-ms time window around the peak, which was typically 3–4 ms after the stimulation artifact. To assess the % silent synapses, only the rates of failures versus successes, not the absolute values of the amplitudes, were used. At +40 mV, successful synaptic responses are conceivably mediated by both AMPARs and NMDARs. Indeed, inhibiting AMPARs by NBQX (5 μM) modestly reduces the amplitudes of EPSCs (Graziane et al., 2016). Despite the effects of NBQX on the amplitudes, the failure rate of synaptic responses at +40 mV is not altered during AMPAR inhibition (Graziane et al., 2016). Thus, in the minimal stimulation assay assessing the % silent synapses, the results are thought not be affected by whether the synaptic responses +40 mV are mediated by NMDARs alone or by both AMPARs and NMDARs.

### DiI staining of spine density and morphology

Rats were perfused with 0.1M PB followed by 1.5% PFA. The brains were dissected out and post-fixed in 1.5% PFA for 1 h before sectioning. Coronal sections (150-µm thick) containing the NAcSh were collected using a Leica VT1200S vibratome, and mounted onto glass slides. A circle around each slide was drawn with a solvent-resistant histology pen, and phosphate buffered saline (PBS) solution was added inside the circle to prevent tissue from drying. DiI fine crystals (Invitrogen) were delivered under a dissecting microscope onto the surface of slices using a brush with stiff hairs. DiI was allowed to diffuse in slices covered by PBS solution for 24 h at 4°C, and then the sections were fixed in 4% PFA at room temperature for 1 h. After a brief wash in PBS, tissues were mounted in aqueous medium (ProLong Gold Antifade, Invitrogen) and sealed with nail polish. Only secondary or tertiary dendrites were included for data acquisition and analysis.

### Immunohistochemistry

The rats were deeply anesthetized with isoflurane, and perfused transcardially first with 10 ml of 10% heparin in 0.1 M phosphate-buffered saline (pH 7.4) and subsequently with 250 ml of 4% paraformaldehyde in 0.1 M phosphate buffer. The brains were post-fixed in the same fixative solution at 4°C for 16 h, followed by graded sucrose concentrations (15% then 30%) in 0.1 M PBS at 4°C for 24 h before sectioning. The brains were sectioned for 45 µm-thick slices at −18°C using Leica Microtome. The brain slices containing the mPFC were collected from +3.2 to 2.8 mm from bregma, and slices containing the NAcSh from +1.8 to 0.8 mm from bregma.

The sections were washed in 0.01 M PBS, then blocked for 1 h in 3% normal donkey serum and 0.3% Triton X-100 in 0.01M PBS. The sections were then incubated with primary antibodies diluted in blocking solution, for 48 h at 4°C with gentle shaking: anti-GFP (1:500, mouse, Abcam, #ab1218), anti-RFP (1:500, rabbit, Rockland, #600-401-379), anti-GFAP (1:500, chicken, Aveslab, #GFAP), anti-NeuN (1:500, rabbit, Cell Signaling Technology, #12941). After incubation with the primary antibody, the sections were rinsed in 0.01 M PBS 6 times, 5 min each time, and were then incubated with the corresponding secondary antibody (donkey anti-mouse Alexa Fluor 488, Abcam, #ab150105; donkey anti-rabbit Alexa Fluor 568, Abcam, #ab175470; goat anti-chicken, Alexa Fluor 488, Abcam, #150169) diluted in the blocking solution for 2 h. After incubation with the secondary antibody, the sections were rinsed in 0.01 M PBS for 6 times, 5 min each time. The sections were then mounted on glass slides, with ProLong Gold Antifade Mountant (ThermoFisher, #P36930), and then secured by coverslip. Fluorescence images were taken with a Leica TCS SP8 confocal laser-scanning microscope, using 10X for low magnification and a 40X oil immersion objective for high magnification. To excite Alexa 488, we used the 488 nm Argon laser with the intensity adjusted to <5% of the maximum output, with the emission filtered between 505-525 nm. Alexa 568 was excited by the 525 nm laser with <10% of the maximum output, with the emission filtered between 590-610 nm.

### Confocal imaging and spine analysis

Images were captured with a Leica TCS SP5 confocal microscope equipped with Leica Application Suite software (Leica). Individually filled neurons were visualized with a 40x oil immersion objective for final verification of their neuronal types (e.g., MSNs vs. interneurons). Individual dendritic segments were identified and scanned at 0.75-µm intervals along the z-axis to obtain a z-stack. After capture, all images were deconvolved within the Leica Application Suite software. Analyses were performed on two-dimensional projection images using ImageJ (NIH). Secondary dendrites were sampled and analyzed due to their significant cellular and behavioral correlates (Graziane et al., 2016). Each experimental groups contained at least 6 animals. For each animal, at least 6 neurons (average 7.6 neurons/animal) were analyzed. For each neuron, at least a 50-µm long dendritic segment was sampled. Similar to our previous studies, we operationally divided spines into 3 categories (Graziane et al., 2016): i) mushroom-like spines were dendritic protrusions with a head diameter >0.5 µm or >2x than the spine neck diameter; ii) stubby spines were dendritic protrusions with no discernable head enlargement and a length of ≤ 0.5 µm; iii) thin/filopodia-like spines were dendritic protrusions with a length of > 0.5 µm and head diameter < 0.5 µm or no discernable head enlargement.

### Astrocytic GCaMP6f imaging in slices

To selectively express GCaMP6f in astrocytes, Aldh1l1-Cre/ERT2 transgenic mice were crossed with ROSA26-Lck-GCaMP6f-flox-GFP transgenic mice (The Jackson Laboratory, 029655 and 029626) to generate offspring carrying both transgenes. The 8-16 week-old double-mutant mice (both male and female) received 5 daily i.p, injections of tamoxifen (100 mg/kg, dissolved in corn oil, VWR, #9001-30-7). The expression of GCaMP6f was verified by immunohistochemistry (with antibodies targeting GFP and neural marker NeuN).

Three weeks after tamoxifen injections, coronal brains slices (260 μm thick) containing the NAcSh were prepared, initially incubated at ∼34°C for 30 min, and subsequently kept at room temperature (∼22°C) in aCSF, saturated with 95% O2 and 5% CO2. Video of GCaMP6f-mediated fluorescent activities in slices were captured by a Hamamatsu ORCE-ER C4742 CCI camera mounted on an Olympus BX51W1 microscope through the 488-nm filter. The camera was set with a fixed exposure time of 200 ms and a frame speed of 5 fps during recording. The temperature of the imaging chamber was set at ∼31°C. The imaging was confined to the medial-ventral portion of the NACSh through a 40x water immersion objective lens. For each video recording, the sampling area of the slice was 3.6 mm^2^ (1344 x 1024 pixels).

The basal GCaMP6f-mediated activities were recorded twice, each lasting 60 sec, separated by a 5 min interval. During this baseline recording, the slices were continuously perfused by aCSF (vehicle control). After establishing the baseline, aCSF that contained 20 µM cocaine hydrochloride was then perfused into the recording chamber. The stock solution of 100 mM cocaine hydrochloride was pre-prepared in aCSF, and was diluted 5000 times upon use, with the osmolarity and pH adjusted to regular levels of aCSF. After 5 min of initial perfusion of cocaine, GCaMP6f activities were recorded twice, with each recording lasting 60 sec, separated by a 5 min interval. The bright field images were checked before and after video recording to detect potential changes in slice position or other imaging conditions, changes of which disqualified the data for further analysis.

The images were analyzed with both ImageJ and Python. The fluorescent signals in the videos were first pre-processed with the rolling ball algorithm in ImageJ to remove uneven background illuminations across frames. Each video was then subjected to motion correction, source extraction, and deconvolution in Python using the Calcium Imaging Analysis (CaImAn) software package (Giovannucci et al., 2019). The fluorescent signals then underwent a high-pass filter and motion-correction using the NoRmCorre algorithm (Pnevmatikakis and Giovannucci, 2017). A modified version of the constrained non-negative matrix factorization framework (CNMF-E) specialized for de-mixing and de-noising one photon data was used to extract fluorescent signals from the videos (Pnevmatikakis et al., 2016; Zhou et al., 2018). The spatial-temporal data (shapes, locations, and peaks of the calcium transients in the field of view) extracted by the CNMF-E were used to identify and classify fluorescent events (Friedrich et al., 2017). To isolate high signal astrocyte activities and reduce noise and neuropil contamination, only fluorescent events with a signal-to-noise ratio above a threshold of 2 were kept for further data analysis. A customized Python code was written to analyze calcium transients. Calcium transients were included in data analysis when their peak heights were at least 2-fold higher than the standard deviation, and at least 1000 ms apart. The calcium transients were plotted by using calcium transients/fluorescent signals extracted by the CNMF-E. The number of locations of calcium-mediated events were determined by the total numbers of region of interests (ROIs) where the extracted fluorescent events had a signal-to-noise ratio at least 2-fold higher than the threshold in the field of view. Each ROI may represent separate astrocytes or separable functional domains within astrocytes. The number of events was determined by the total numbers of calcium transient events during the entire periods of recording. The frequency (events/min) was determined by averaging the number of calcium transients per min within the field of view. The areas of transients were determined by the total net changes in fluorescent signals of calcium transients over the baseline multiplied by the duration of the transients. The mean amplitudes were determined by the mean amplitudes of all the calcium transients of all the ROIs. The averaged peak amplitudes were determined by the average of the maximum calcium transients of all the ROIs.

### Drugs and reagents

Chemicals were purchased from Tocris, Sigma-Aldrich. ThermoFisher, J.T.Baker, Acros, Fluka, Toronto Research Chemicals, Cayman Chemicals, and Spectrum. All reagents administrated into animals were purchased from Henry Schein, Zoetis, Putney, Vedco, Sagent, and Bimeda. For viruses, the AAV2/5-gfaABC1D-tdTomato was purchased from AddGene (44332), and the AAV2/5-hSyn-hChR2(H134R)-mCherry was purchased from the UNC Vector Center. The plasmids were purchased from AddGene, including pZac2.1-GfaABC1D-mCherry-hPMCA2w/b (111568) and pcDNA3 mTSP2 (12411).

### Data acquisition and statistics

All results are shown as mean ± s.e.m. All experiments were replicated in 5-16 rats. All data collection was randomized. All data were assumed to be normally distributed, but this was not formally tested. No statistical methods were used to pre-determine sample sizes, but our sample sizes are similar to those reported in previous publications (Graziane et al., 2016; Huang et al., 2009; Lee et al., 2013; Ma et al., 2014; Neumann et al., 2016; Wright et al., 2019). All data were analyzed offline and investigators were blinded to experimental conditions during the analyses.

A total of 468 rats and 147 mice were used for this study, among which 170 rats and 33 mice were excluded from the final data analysis and interpretation due to the following reasons: 1) 76 rats were excluded because of health issues after surgeries (e.g., >20% drop in body weight in a day), failed catheterization, or failure to reach the self-administration criteria; 2) 8 rats were excluded due to off-target stereotaxic injections or poor viral expression; and 3) 86 rats and 33 mice were excluded because of experimental failures (e.g., unsuccessful slice preparations, failed recordings, or other experimental incidents). No animals were excluded after data acquisition was accomplished.

Repeated experiments for the same group were pooled together for statistical analysis. Sample sizes were based on our previous studies performing similar experiments (Graziane et al., 2016; Huang et al., 2009; Lee et al., 2013; Ma et al., 2014; Neumann et al., 2016; Wright et al., 2019). For electrophysiology and dendritic spine experiments, the sample sizes are presented as n/m, where n = number of cells and m = number of rats. For behavioral experiments, the sample sizes are presented as n, which is the number of rats. Animal-based statistics were used for all data analyses. For electrophysiological experiments, we averaged the values of all the cells from each rat to obtain an animal-based mean value for statistical analysis (Graziane et al., 2016; Wright et al., 2019). For dendritic spine experiments, we averaged individual dendritic segment values from each cell to obtain a cell-based value, and then averaged cell-based values from each rat to obtain animal-based values for statistical analysis(Graziane et al., 2016; Wright et al., 2019). Statistical significance was assessed using two-tailed unpaired t-tests, one-way or two-way ANOVA, or mixed two-way ANOVA with repeated measure followed by Bonferroni posttest, or repeated measures two-way ANOVA, as specified in the related text. Two-tailed tests were performed for analyses. Statistical significance was set at *p* < 0.05 for all experiments. Statistical analyses were performed in GraphPad Prism (v7) and SPSS v19 (IBM).

## Acknowledgement

We thank the late Dr. Ben Barres for insightful suggestions on key experimental designs, Dr, Mary Torregrossa and Yavin Shaham for suggestions on behavioral experiments, Jaryd Ross and Kevin Tang for technical support. The study was supported by NIH NIDA DA029565, DA028020, DA023206 and DA024570, DA031551, NS107604(O.M.S) the Göttingen Graduate School for Neurosciences, Biophysics, and Molecular Biosciences [Grant GSC226/1] (to AB and AS). Cocaine was provided by the NIH NIDA drug supply program.

---

**Figure S1.**
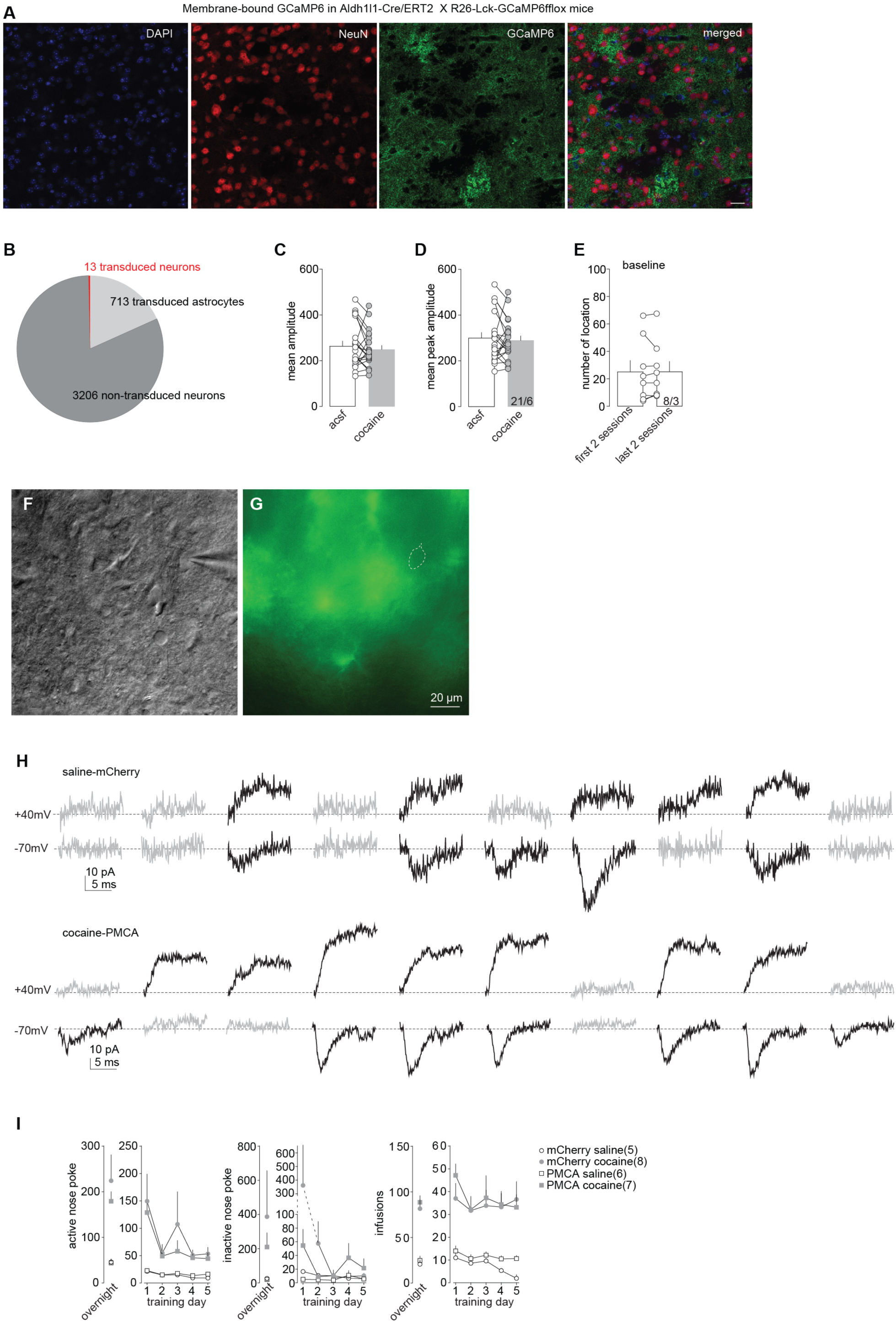
Supplementary materials for. **Figure 1. A** The same images presented in Figure 1B-C with additional DAPI histochemical staining and merged images of triple staining. **B** Summary of astrocyte-specific expression. The results were analyzed by including both the astrocytic expression of GCaMP6f using the Aldh1/1-Cre/Ert2 x Rosa26-Lck-GCaMP6f-flox mice and AAV-mediated astrocyte-selective expression using the hPMCA2w/b promoter. **C**, **D** Summaries showing that neither the mean amplitudes (t_1,20_ = 0.86, p = 0.40; **C**) nor the maximum amplitudes (t_1,20_ = 0.54, p = 0.60; **D**) of astrocytic calcium transients were altered by perfusion of cocaine. **E** Summary showing that the total calcium locations that exhibited calcium-mediated events remained the same throughout the experimental procedure (t_1,7_ = 0.07, p = 0.94). **F,G** Example DIC (**F**) and fluorescent (**G**) images showing the recorded NAcSh MSNs were located within the astrocytic matrix. **H** Example EPSCs from the minimal stimulation assay of silent synapses showing clear distinction between successes and failures. These two sets of EPSCs are the same as presented in Figure 1T and 1W but are spread out for clarity. **I** Summaries of self-administration results of rats presented in Figure 1T-X.

**Figure S2.**
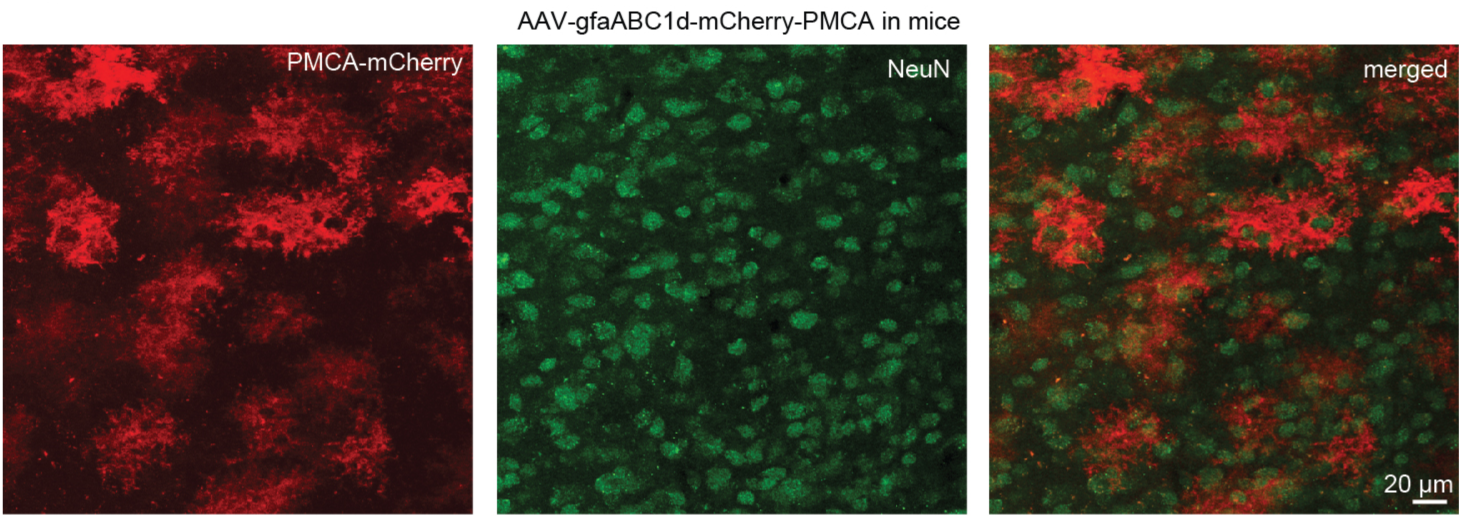
GfaABC_1_-D AAV-mediated expression of hPMCA2w/b-mCherry in mice. Example confocal images of a mouse NAcSh slice with GfaABC_1_D AAV-mediated expression of hPMCA2w/b-mCherry, in which mCherry signals depicted bushy, astrocytic morphologies, with minimal overlap with NeuN signals (green).

**Figure S3.**
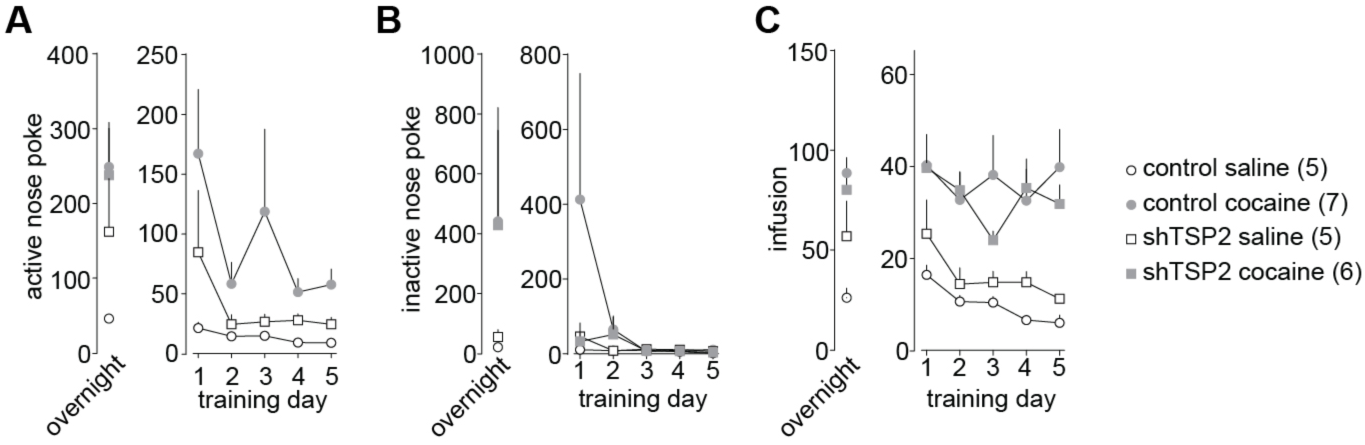
Supplementary materials for. **Figure 3. A-C** Summaries of self-administration results from rats presented in Figure 3J-N.

**Figure S4.**
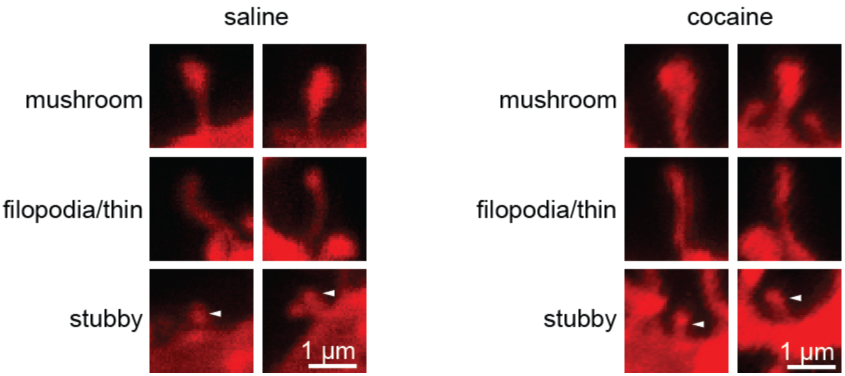
Supplementary materials for. **Figure 2**. Examples of mushroom-like, filopodia-like, long-thin, and stubby spines from the results presented in Figure 4G-L.

**Figure S5.**
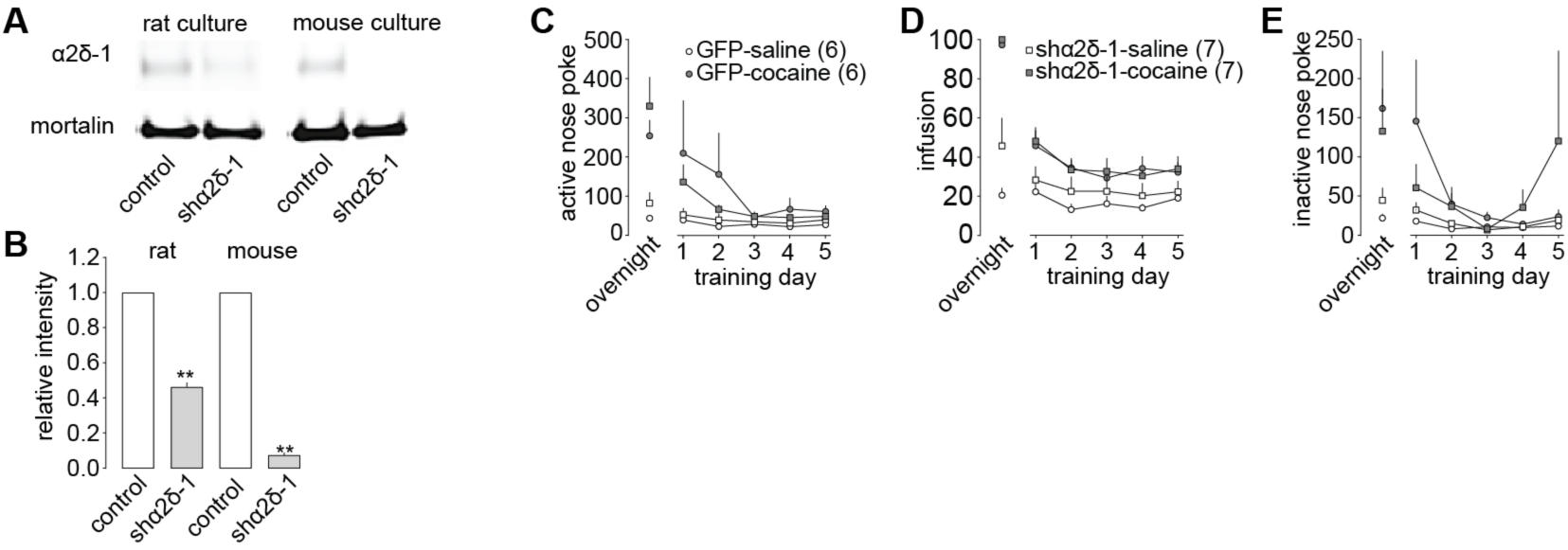
Supplementary materials for. **Figure 5. A,B** Western analyses showing the knockdown efficacies of shα2δ-1 in neuronal cultures (rats: t_1,2_ = 37.79, p < 0.01; mice: t_1,2_ = 112.86, p<0.01, paired t-test). **C-E** Summaries of self-administration results from rats presented in Figure 5E-I.

**Figure S6.**
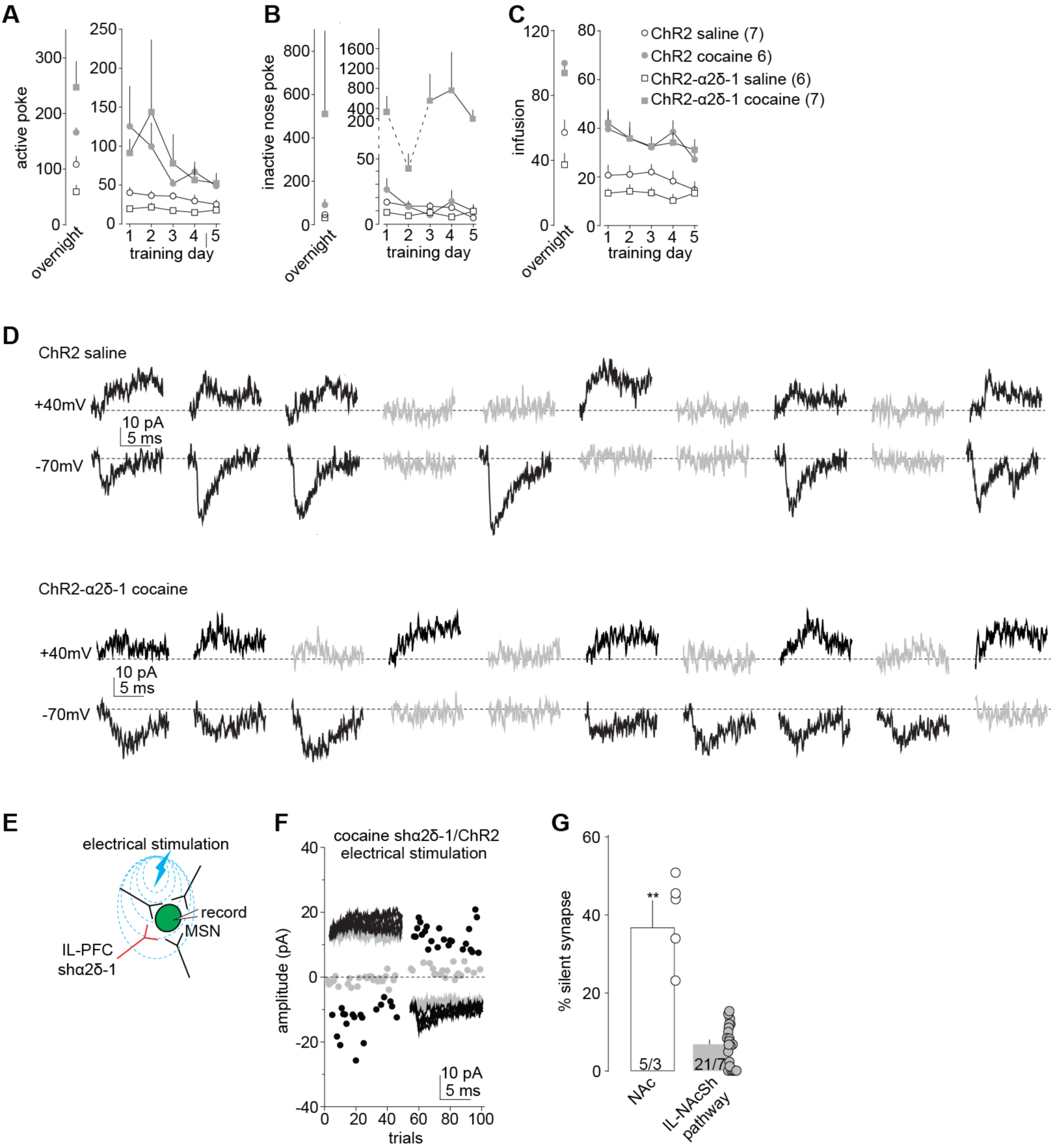
Supplementary materials for. **Figure 6. A-C** Summaries of self-administration results from rats presented in Figure 6H-L. **D** Example EPSCs from the optogenetic minimal stimulation assay of silent synapses showing clear distinction between successes and failures. These two sets of EPSCs are the same as presented in Figure 6H and 6K but are spread out for clarity. **E** Diagram showing that the minimal stimulation assay by electrical stimulation presumably also sampled presynaptic terminals that did not express shα2δ-1 and ChR2. **F, G** Example trials (**F**) and summary (**G**) showing that among non-selectively sampled synapses, the increased % silent synapses was detected in cocaine-trained rats with PFC expression of shα2δ-1 and ChR2 (t_1,8_= 8.99, p < 0.01), while this increase was prevented in the IL-to-NAcSh projection in which presynaptic terminals expressed shα2δ-1 and ChR2. Note that the data for IL-to-NAcSh projection are the same as presented in Figure 6L.

**Figure S7.**
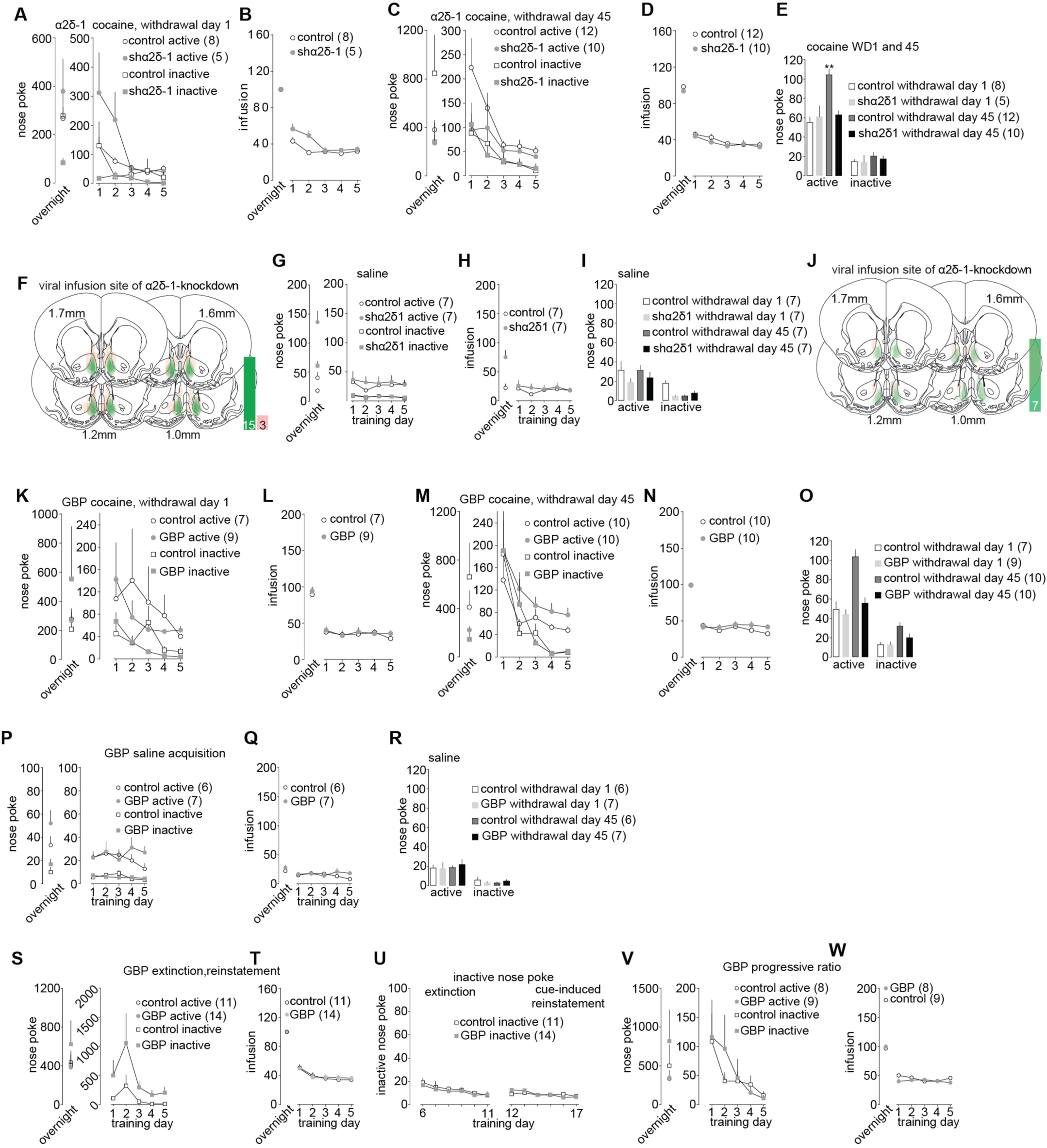
Supplementary materials for. **Figure 7. A,B** Cocaine self-administration results for rats that were tested on withdrawal day 1. These rats were used as controls for rats tested on withdrawal day 45, which were presented in Figure 7A-D. **C.D** Additional self-administration results from rats presented in Figure 7A-D. **E** Summary of cue-induced cocaine seeking in GFP and shα2δ-1 rats on withdrawal days 1 versus 45. GFP control rats exhibited incubation of cue-induced cocaine craving and this phenomenon was prevented in shα2δ-1 rats. Data for withdrawal day 45 were the same as presented in Figure 7D. **F** Injection sites of AAV in rats presented in Figure 7B-D. **G-I** Results of saline self-administration in rats used as controls for cocaine-trained rats presented in Figure B-D. **J** Injection sites of AAVs is saline control rats. **K-L** Cocaine self-administration results for rats that were tested on withdrawal day 1. These rats were used as control for rats tested on withdrawal day 45, which were presented in Figure 7E-H. **M,N** Additional self-administration results from rats presented in Figure 7E-H. **O** Summary of cue-induced cocaine seeking in vehicle and GBP rats on withdrawal days 1 versus 45. Vehicle rats exhibited incubation of cue-induced cocaine craving, and this phenomenon was prevented in GBP rats. **P-R** Results of saline self-administration in rats used as controls for cocaine-trained rats presented in Figure 7E-H. **S-U** Additional results related to self-administration, extinction, and reinstatement from rats presented in Figure 7I,J. **V,W** Additional self-administration results from rats presented in Figure 7K.

